# Divergent granulopoiesis at extramedullary sites safeguards host defense

**DOI:** 10.1101/2025.02.25.638781

**Authors:** Carlos Silvestre-Roig, Raphael Chevre, Alexander Bender, Lina M. Vöcking, Ali Hageb, Mathis Richter, Merieme Farjia, Quinte Braster, Mauricio Guzman, Jordi Sintes, Samriti Sharma, Patricia Lemnitzer, David Ahern, Collins Osei-Sarpong, Daniel R. Engel, Claudia Monaco, Petra Dersch, Artur Kibler, Andrea Cerutti, Triantafyllos Chavakis, Jadwiga Jablonska, Oliver Soehnlein

**Author notes:** Correspondence should be addressed to CSR or OS or.

## Abstract

Extramedullary organs such as the spleen can assume granulopoiesis as a supportive mechanism to cope with the demands during persistent inflammation. However, the quantitative output of extramedullary granulopoiesis is limited, thus raising the question if the spleen in fact provides neutrophils of a qualitative difference rather than merely contributing to neutrophil numbers. Here we report splenic stress granulopoiesis with distinct production and differentiation trajectories. Myeloid progenitors in the spleen engage in accelerated production of neutrophils with an immature phenotype. Yet, neutrophils generated during persistent stress granulopoiesis are fully competent to exert antimicrobial functions and are necessary to contain bacterial invasion. Activation of type I interferon signaling in the spleen is required for splenic neutrophil production and its loss impairs host defense. Thus, the spleen provides an immunological environment for stress-induced rapid production and priming of highly active neutrophils to meet the demands during infection.

## Introduction

Neutrophils are the major constituent of the innate arm of the host response against invading pathogens. Production and mobilization of neutrophils are tightly regulated processes that control the daily level of circulating neutrophils during steady state (Yvan-Charvet and Ng, 2019). However, this precise regulation is dramatically altered during acute inflammation leading to a massive discharge of neutrophils. Persistence of the inflammatory signals in not rapidly cleared infections initiates a program of emergency granulopoiesis facilitating the recovery of the emptied neutrophil bone marrow pool and maintaining the supply of neutrophils in the circulation with distinct levels of maturation (Swann et al., 2024; Yvan-Charvet and Ng, 2019). Under such conditions, hematopoietic stem and progenitor cells (HSPCs) and myeloid progenitors can be mobilized to the circulation and engraft in secondary organs such as the spleen or liver in a process termed extramedullary hematopoiesis (EMH) (Mende and Laurenti, 2021). In mice, this process occurs not only during infection (Burberry et al., 2014; Dutta et al., 2015) but extends to other states of sustained inflammation such as cardiovascular disease (Leuschner et al., 2012; Robbins et al., 2012) or cancer (Wu et al., 2018). Similarly, counts of circulating hematopoietic stem cells (HSCs) and progenitors increase in individuals suffering from such pathologies. EMH permits outsourcing neutrophil production to secondary organs for the bone marrow to compensate for a deficit in physical space; however, the limited neutrophil output by extramedullary production is at odds with this concept.

Neutrophil functionality can be shaped across their production and differentiation pipeline (Xie et al., 2020), a process that can be beneficial for the host upon inflammatory pressure. Thus, we here hypothesize that stress-induced granulopoiesis independent of the bone marrow arises as a host mechanism to meet qualitative rather than quantitative neutrophil output in response to lasting inflammatory insults. Here, we investigate the importance of extramedullary granulopoiesis as a supportive mechanism for neutrophil production under inflammatory stress. Using epigenomic and functional profiling of neutrophils across the differentiation tree in the bone marrow and spleen, we show that the spleen serves as a site that rapidly produces neutrophils with enhanced antimicrobial function. We demonstrate that type I interferon signaling primes neutrophil progenitors for express differentiation into neutrophils which are prematurely released into the circulation. Functional analysis demonstrated that this premature subpopulation exhibits fully competent effector functions facilitating microbe elimination.

## Results

### Alternative granulopoiesis emerges from extramedullary production

To comprehensively profile emergency granulopoiesis in the spleen, we mapped the hematopoietic differentiation dynamics from hematopoietic stem and progenitor cells to differentiated immature and mature neutrophils across chronic conditions that alter neutrophil production by flow cytometry (**Figure 1, Extended Figure 1**). C57BL6/J mice subjected to models of melanoma, repeated lipopolysaccharide (LPS) administration, aging, and hypercholesterolemia were profiled from Lin^-^Sca1^+^cKit^+^ (LSKs) cells all the way to immature (immNEU) and mature (matNEU) neutrophils within the bone marrow and the spleen (**Figure 1A/B, Extended Figure 1B-G**). Splenic myelopoiesis generated a strong neutrophil-biased hematopoiesis with a striking expansion of preNEUs in absolute counts and the differentiation to immNEU and matNEU respective to spleen-resident cells found in steady-state. This expansion of neutrophil progenitors and descendants in the spleen mimicked bone marrow granulopoiesis under steady-state (**Figure 1A, Extended Figure 1D**); importantly, granulopoiesis in the bone marrow was only mildly affected during stress conditions studied here (**Figure 1B, Extended Figure 1D**). Our results suggest that - under a variety of persistent challenges - the spleen engages in active granulopoiesis with expansion of neutrophil progenitors and elevated production of neutrophils peaking at the immature stage. To simplify and harmonize these models we made use of a model of myeloablation (Inra et al., 2015) based on treatment with one single dose of cyclophosphamide (CPM) followed by repeated administration of the CXCR4 antagonist (AMD3100, model of CPM/AMD); this model recapitulated the commonality of stress-induced granulopoiesis in the spleen in all other models with similar proportions and numbers (**Figure 1C, 1D, Extended Figure 1E-G**) while omitting the inflammatory component associated to these. We hence focused on this reductionist model to detail neutrophil production and differentiation at single-cell resolution. Spectral flow cytometry analysis using an antibody panel of 19 markers was used to generate high-dimensional analysis of neutrophil differentiation dynamics employing the PHATE algorithm (Moon et al., 2019) (**Figure 1E**). These analyses confirmed that splenic granulopoiesis is associated with a strong expansion of neutrophil progenitors and immature neutrophils and revealed elevated blood counts of immNEU in these mice. To further detail splenic granulopoiesis at a transcriptomic level, we performed scRNA-seq profiling of myeloid progenitors, neutrophil-committed progenitors (proNEU-preNEU), immature and mature neutrophils. Gr1^+^CD115^+^ (including both neutrophils and monocytes) and cKit^+^ cells from blood, bone marrow and spleen in control or CPM/AMD-treated mice were FACS-sorted (**Extended Figure 2A**) and processed for scRNA-seq analysis. After quality control, a total of 14,042 cells were used for analysis. We next performed unbiased dimensionality reduction and unsupervised clustering on an aggregate of blood, bone marrow and splenic cells from both control and CPM/AMD treatment (**Extended Figure 2B**). Cell annotation using published cell-specific marker datasets and data curation based on differential expression analysis among identified clusters revealed populations of myeloid progenitors, neutrophil-committed progenitors, immature and mature neutrophils, monocytes, NK-cells, and a subpopulation of dendritic-like cells (**Extended Figure 2B**, lower panel). Analysis of cell proportions recapitulated the results observed in our flow cytometry experiments, *i.e.*, a significant expansion of the neutrophil compartment at the expense of monocytic lineages (**Extended Figure 2C**). We next clustered neutrophil progenitors and differentiated neutrophils to analyze neutrophil differentiation pathways in both bone marrow and spleen and their end-stage in the blood (**Extended Figure 2D, 2E**). Analysis of neutrophil subpopulations showed that mature neutrophils dominated the blood compartment in steady-state conditions while immature neutrophils emerged upon EMH (**Figure 1F**) thus confirming our flow cytometry analyses (**Figure 1E**). Interestingly, while the bone marrow remained largely unaltered during EMH, a marked increase in immature neutrophils was found in the spleen. Intriguingly, while the differentiation from the immature to mature neutrophils was continuous in the bone marrow, it was interrupted in the spleen suggesting alterations in the differentiation path and a potential release to the circulation at a premature stage (arrows in **Figures 1E, F**). Importantly, such discontinuous granulopoiesis was also observed in other chronic stress models such as in hypercholesterolemia (**Extended Figure 2F**) and during tumor development (**Extended Figure 2G**). Notably, a significant increase in this subpopulation of immature neutrophils is observed in the blood (**Figure 1E, F**), thus suggesting a rapid mobilization from their splenic niche. Collectively, while minor changes are observed in the bone marrow, the spleen assumes an alternative and interrupted granulopoiesis under inflammatory stress to produce neutrophils whose differentiation stops at a premature stage before being mobilized to the circulation.

**Figure 1.**
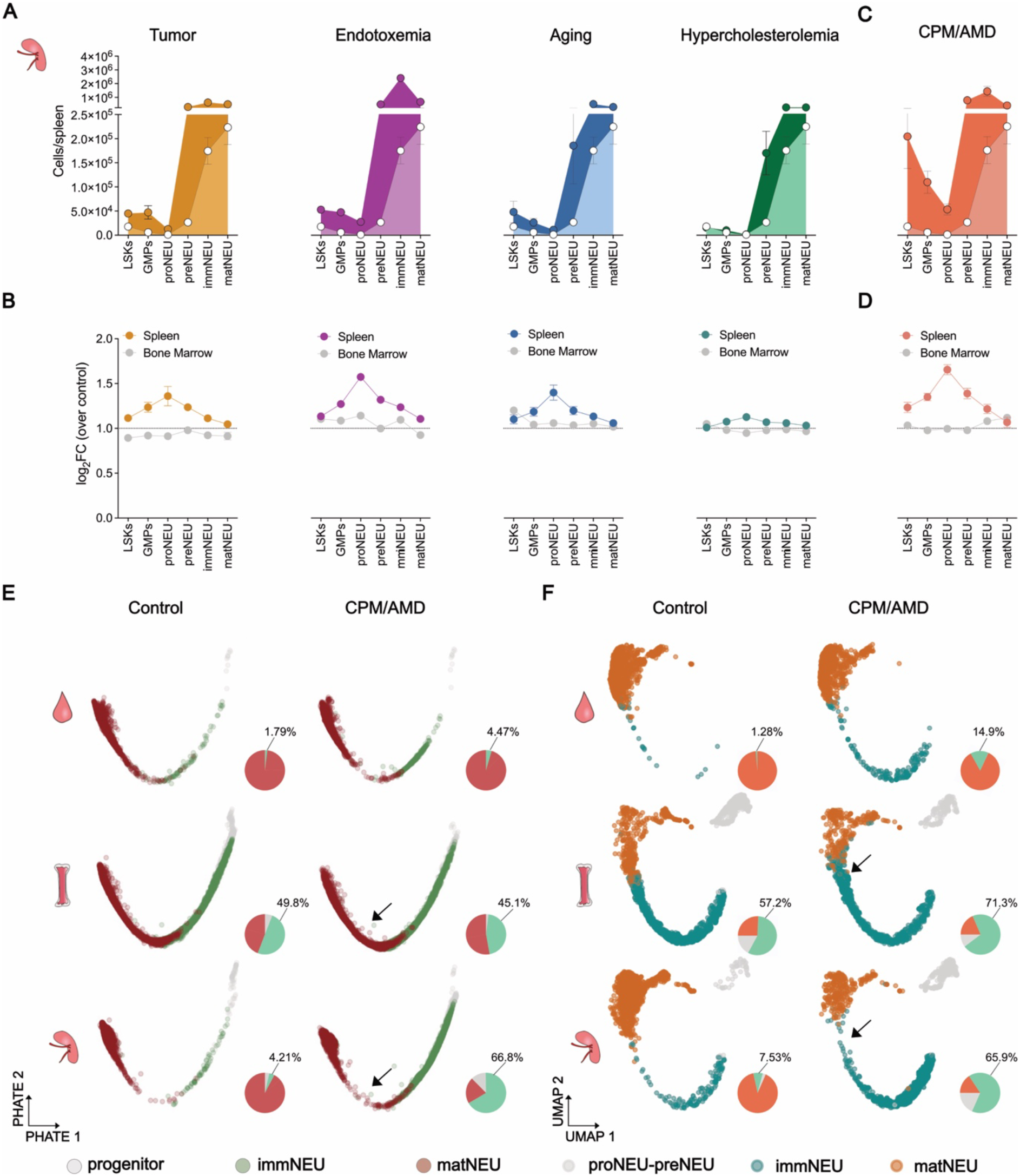
Alternative neutrophil production dynamics in stress-induced splenic granulopoiesis. (A/B) Flow cytometry-based quantification of indicated stages of granulopoiesis in the spleen. LSKs (Lin^-^Sca1^+^cKit^+^), granulocyte-macrophage progenitors (GMP, Lin^-^Sca1^-^cKit^+^CD16/32^high^CD34^+^), neutrophil-specified (proNEUs, Lin^-^Sca1^-^cKit^+^ CD16/32^high^Ly6C^+^CD115^-^CD81^+^CD11b^+^) and-committed progenitors (preNEU, Lin^-^cKit^int^Ly6C^+^CD115^-^CD11b^+^CXCR4^+^), immature (immNEU, CD11b^+^Ly6G^+^CD115^-^CXCR2^-^) and mature neutrophils (matNEU, CD11b^+^Ly6G^+^CD115^-^CXCR2^+^). Data points indicate the mean of 3-10 mice. Lin: Lineage. 8-12 weeks old female C57BL6/J mice with the exception of ageing model. From left to right: tumor model of melanoma, chronic endotoxemia, aging, hypercholesterolemia. (A) Empty circles indicate respective splenic cell populations under steady-state (control) condition. Filled circles represent counts for each maturation stage in each model. (B) Log2 fold change of each indicated stages over respective control in bone marrow or spleen granulopoiesis. **(C/D)** C57BL6/J female mice (8-12 weeks old) were administered with cyclophosphamide (CPM), followed by daily administration of 5 mg/kg of AMD3100 (AMD). Flow cytometry-based enumeration (C) and log2 fold change over respective control (D) of stages of granulopoiesis in bone marrow and spleen. **(E)** PHATE analysis of flow cytometry data on neutrophils and their progenitors from control or CPM/AMD-treated mice of indicated organs. Pie charts depict the proportion of neutrophil progenitors, immature (immNEU) and mature (matNEU) neutrophils. Percentage for immNEU is indicated. n = 5 mice. **(F)** Uniform manifold approximation and projection (UMAP) plot of aggregated cells from scRNA-seq analysis of blood, bone marrow and spleen neutrophil progenitors and descendants colored by cell type. Pie charts depict the proportion of neutrophil progenitors (proNEU_preNEU), immature (immNEU) and mature (matNEU). Percentage for immNEU is displayed. n = 4 mice. Arrows point at continuous vs disrupted granulopoiesis in bone marrow or spleen, respectively.

### Splenic granulopoiesis provides the circulation with immature neutrophils

We next sought to determine the contribution of splenic granulopoiesis to circulating neutrophil populations. Herein, we performed mass cytometry analysis using a panel of 35 markers on blood, bone marrow, and spleen leukocytes in vehicle and CPM/AMD-treated mice (**Figure 2A**). After normalization, high-dimensionality reduction and unbiased clustering, we annotated the leukocytes based on their expression of lineage markers (**Extended Figure 3A**). Next, we focused on Ly6G^+^ neutrophils and performed unbiased clustering on selected cells, identifying 15 metaclusters (**Figure 2A, Extended Figure 3B**) with differential enrichment across organs and conditions (**Extended Figure 3C**). Based on their relative abundance by organ (**Figure 2B**) and expression of surface markers (**Figure 2C**), we identified marrow (NeuP) and splenic (sNeuP) neutrophil progenitors, marrow immNEU (Neu1) and matNEU (Neu2), splenic (sNeu) and circulating (Neu3) neutrophils. Subpopulations with a lower abundance (< 3% of all metaclusters) were not annotated. When focusing on clusters emerging in the spleen during EMH, we found that these subpopulations appeared in the circulation during stress-induced granulopoiesis while not changing in the bone marrow compartment, suggesting a splenic origin (**Figure 2D**). To validate these results, we studied the dynamics of neutrophil differentiation and mobilization to the circulation using 5’ bromo-deoxy-uridine (BrdU) pulse-chase experiments (**Figure 2E**). Mice were injected with BrdU 3 days before sacrifice, time required for the BrdU signal to label dividing neutrophil progenitors and to reach the post-mitotic pool (immature and mature neutrophils) before being released to the circulation. We then evaluated the BrdU signal across granulopoiesis as defined by PHATE trajectory (Moon et al., 2019) at Zeitgeber time (ZT) 13, *i.e.* the time of neutrophil release to the circulation (Casanova-Acebes et al., 2013), and at ZT2 (day time) (**Figure 2E**). Interestingly, BrdU^+^ neutrophils were found to progress with time within the bone marrow from an immature (PHATE^low^) to a mature (PHATE^high^) state while remaining immature in the spleen. We then binned cells across PHATE dimension 1 into 8 equal-sized clusters (**Figure 2F)**. These clusters defined cells with progressive maturation levels (from bin 1 to 8) as demonstrated by CXCR2 expression (**Figure 2G**). Analysis of the proportion of these binned clusters on BrdU^+^ neutrophils showed an absent increase in bin 8 within the spleen thus supporting the inability of immature splenic neutrophils to mature (**Figure 2H**). These results suggested the possibility that splenic neutrophils are prematurely released to the circulation at an immature state constituting approximately 5-10% of the total of circulating neutrophils in mice treated with CPM/AMD, fed high-fat diet, or repeatedly injected with LPS (**Extended Figure 3D-F**). Furthermore, these results also suggest the disconnection between splenic immature neutrophils and their mature counterparts. Indeed, Pearson correlation analysis of mature and immature neutrophils shows a strong correlation in the bone marrow (**Extended Figure 3G**), but not in the splenic niche (**Extended Figure 3H**). These results would, therefore, indicate that mature neutrophils in the spleen are likely recruited from the circulation rather than being produced *in situ*. Consistently, we found in additional BrdU pulse-chase experiments that while immature neutrophils progressively acquired the signal, mature neutrophils positive for BrdU appeared in the spleen only at the time of their peak in the blood (**Extended Figure 3I**). Altogether, splenic neutrophils constitute a mixture of *de novo*-produced immature neutrophils and mature blood-borne neutrophils.

**Figure 2.**
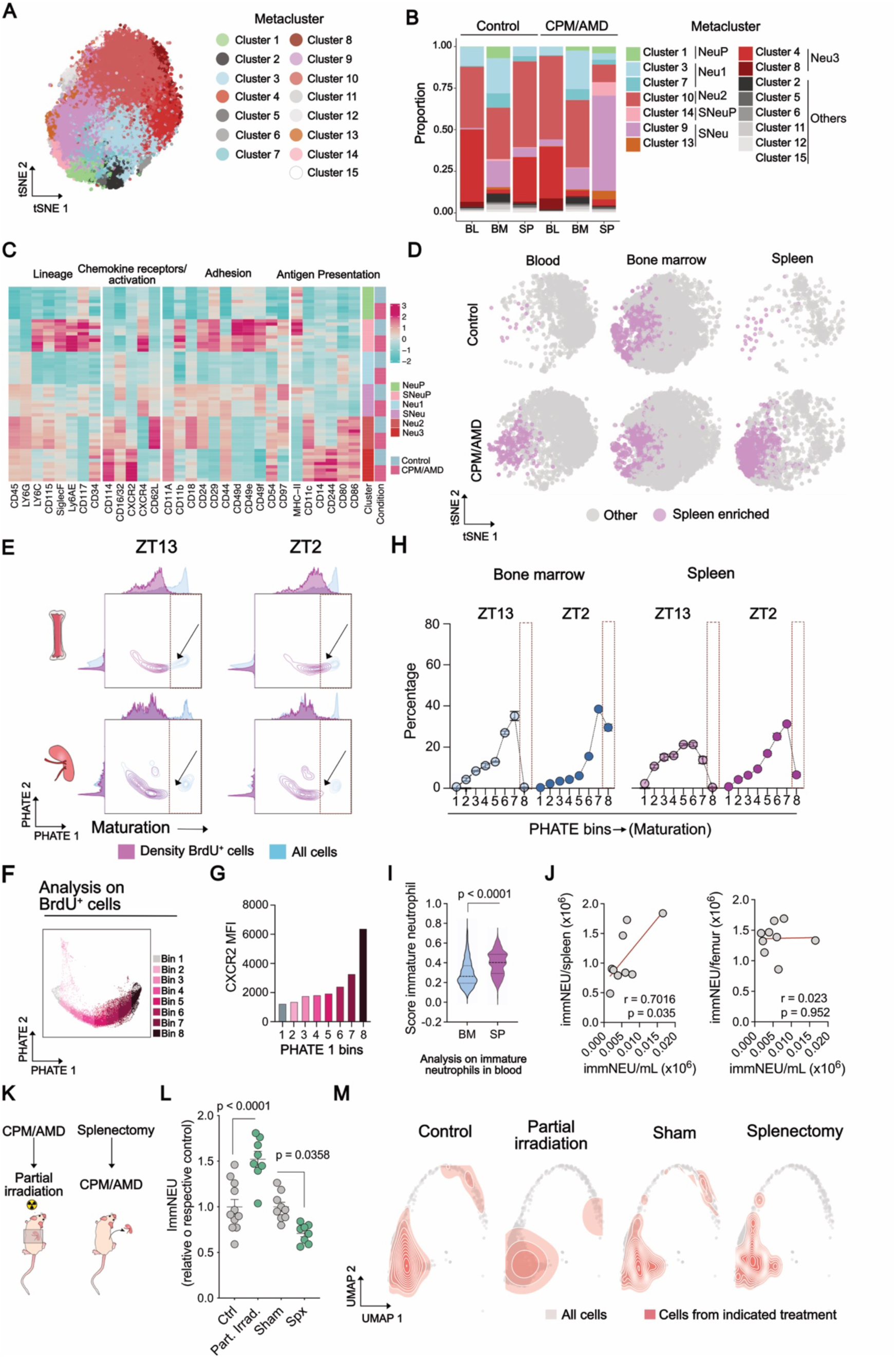
**Premature release of splenic immature neutrophils contributes to the blood neutrophil pool**. C57BL6/J female mice were administered with cyclophosphamide (CPM), followed by daily administration of 5 mg/kg of AMD3100 (AMD). **(A-D)** Mass cytometry analysis of blood, bone marrow and spleen neutrophils in steady state or after CPM/AMD treatment. n=5 mice/condition. **(A)** tSNE plot of aggregated cells from blood, bone marrow and spleen neutrophil progenitors and descendants colored by metacluster. (**B**) Proportion analysis of enriched subpopulations within blood, bone marrow and spleen under Vehicle (Control) or CPM/AMD-treatment. BL: Blood; BM: Bone marrow; SP: Spleen. (**C**) Heatmap showing scaled expression of surface receptors within each enriched subpopulation, grouped by function. (**D**) tSNE plot showing spleen-enriched population (magenta) over all cells (light gray). **(E-H)** Mice receiving CPM/AMD were injected 5‘Bromo-deoxyuridine (BrdU) 3 days before sacrifice at Zeitgeber Time (ZT)13 or ZT2. **(E)** Plots showing PHATE analysis in indicated organs and time points. Blue and magenta density depicts all and BrdU^+^ lineage negative cells (Lin: B220^+^CD19^+^CD90.2^+^CD3^+^NK1.1^+^F4/80^+^ FcRε^+^TER119^+^), respectively. Dashed line indicates the disruption in maturation, while the arrow points at the progression of maturation in the bone marrow. **(F)** Binning of PHATE1 dimension. **(G)** CXCR2 expression across PHATE1 bins. **(H)** Percentage of PHATE1 bins within BrdU^+^ neutrophils for indicated organs in CPM/AMD-treated mice. Dots indicate mean of 5 mice. **(I)** Enrichment analysis of bone marrow or spleen immature signatures obtained from scRNA dataset assessed in circulating immature neutrophils. **(J)** Pearson correlation of immature neutrophils in bone marrow vs blood (left panel) and spleen vs blood (right panel). Every dot represents one mouse. **(K-M)** Mice under CPM/AMD were subjected to splenectomy or partial irradiation shielding the spleen. **(K)** Scheme of experimental procedure. **(L)** Quantification of immature neutrophils (Ly6G^+^CD101^low^CXCR2^low^) in blood. Data are mean±SEM, two-sided t-test analysis relative to the mean of the respective control. **(M)** scRNA-seq analysis of circulating neutrophils from mice subjected to splenectomy or sham followed by CPM/AMD treatment or mice subjected to CPM/AMD treatment followed by partial irradiation with 6.5 Gy combined with lead radiation shielding of the abdominal area. UMAP plot showing aggregated cells from all treatments (grey) and overlayed density of cells from indicated treatment (red).

In line with the concept of a splenic origin of immature neutrophils in the blood, this subpopulation scored higher in our scRNA-seq data for a transcriptomic signature of splenic immature neutrophils as compared to bone marrow immature neutrophils (**Figure 2I**). In further support of this notion, circulating immature neutrophil counts positively correlated with immature neutrophils in the spleen but not the bone marrow (**Figure 2J**). Moreover, pairwise analysis showed minimal differences between sNeu and immature blood neutrophils (**Extended Figure 3J**). Of note, splenic or blood mature neutrophils showed significant transcriptional differences respective to their immature counterparts. These differences were minimal when comparing splenic mature neutrophils to circulating mature cells (**Extended Figure 3J**), suggesting that mature splenic neutrophils are primarily blood-derived as reported previously (Ballesteros et al., 2020; Casanova-Acebes et al., 2018). To causally prove the contribution of the splenic granulopoiesis to the circulating neutrophil pool, we performed splenectomy or partial irradiation shielding the abdomen to protect active splenic granulopoiesis prior or after induction of emergency granulopoiesis, respectively (**Figure 2K**). As predicted, flow cytometry analysis (**Figure 2L**) or scRNA-seq analysis (**Figure 2M**) of circulating neutrophils revealed that splenectomized mice showed a reduced presence of immature neutrophils; partial irradiation, on the other hand, led to enrichment of circulating immature neutrophils. In summary, splenic granulopoiesis generates neutrophils that prematurely exit their site of production delivering immature neutrophils to the circulating neutrophil pool.

### Clustered myeloid progenitors engage in accelerated splenic granulopoiesis

Stress-induced granulopoiesis copes with neutrophil demand by conveying a rapid stem cell division and enhanced cell differentiation towards neutrophils. Mechanistically, under stress conditions, neutrophil differentiation is fast-tracked by shorter differentiation pathways. This is facilitated by clustered myeloid progenitor positioning close to vascular niches (Herault et al., 2017) and enhanced abundance of growth factors such as G-CSF (Yvan-Charvet and Ng, 2019). We hence speculated that sNeuP engage in a rapid differentiation process preceding the premature release of neutrophils to the circulation thus offering an explanation for the enrichment of an immature stage. BrdU pulse experiments performed within a 24-hour time window showed a profound dilution of the signal from the preNEU towards the immNEU state within the spleen as compared to the bone marrow (**Figure 3A, Extended Figure 4A**), suggesting an accelerated maturation at the preNEU to immNEU intersect. Furthermore, splenic preNEUs were intrinsically poised for division and differentiation, as isolated preNEUs incubated with growth factors or bone marrow supernatants gave rise to an increased number of descendants (**Extended Figure 4B**). To test the importance of the splenic environment to activate these progenitors, bone marrow-isolated progenitors from dsRED mice (red-fluorescent) were adoptively transferred into the spleen of C57BL6 mice by subcapsular injection (**Figure 3B, Extended Figure 4C**). Eight days after the injection, splenic progenitors differentiated into neutrophils, and we observed a positive correlation between cKit^+^ cells and neutrophils in the spleen (**Extended Figure 4C**). Interestingly, dsRED^+^ neutrophils exhibited a reduced maturation (lower expression of CXCR2) as compared to endogenous neutrophils (**Figure 3B**), suggesting that the spleen environment was sufficient to drive this altered cell differentiation.

**Figure 3.**
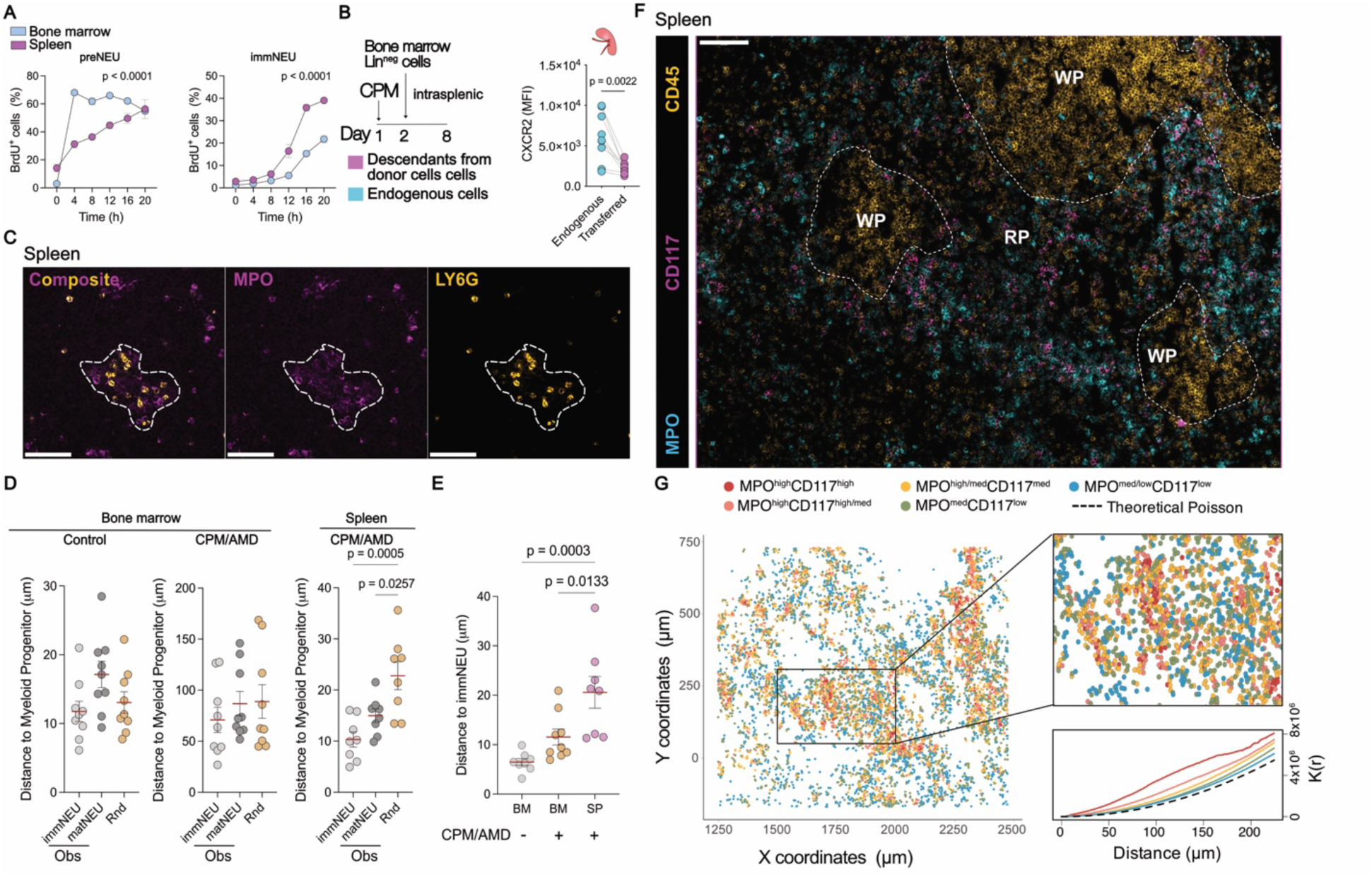
**Accelerated differentiation of splenic neutrophils from spatially clustered progenitors**. **(A)** CPM/AMD-treated mice were injected with a pulse of 5‘Bromo-deoxyuridine (BrdU) and sacrificed at indicated time-points. Shown is the percentage of BrdU^+^ cells. Data are mean±SEM. Two-way ANOVA with Šidák’s post-hoc correction. n = 2-4 mice per group. **(B)** Mice were subjected to CPM followed by subcapsular splenic injection of 2×10^5^ lineage negative cells and sacrificed 8 days after. CXCR2 expression in donor respective to endogenous neutrophils in the spleen. **(C/D)** Immunofluorescence images depicting nuclei (DAPI, blue), Myeloperoxidase (MPO, magenta), Ly6G (ochre) in the red pulp of the spleen of CPM/AMD-treated mice. Scale bar: 100 μm. Neutrophil production cluster delineated by dashed white line. **(D/E)** Distance analysis of indicated cell subpopulations. (D) Analysis of immature (MPO^med^Ly6G^med^) and mature (MPO^low^Ly6G^high^) neutrophils to myeloid progenitors (MPO^high^Ly6G^low^). Obs: Observed; Rnd: Random. One-way ANOVA with Tukey’s post-hoc correction. n = 8-9 images from 3 mice/group. Data are mean +/-SEM. (**E**) Quantification of distance (pixels) of mature vs immature neutrophil. One-way ANOVA with Tukey’s post-hoc correction. n = 8-9 images from 3 mice/group. Data are mean +/-SEM. (**F**) Immunofluorescence images depicting MPO (cyan), CD117 (magenta) and CD45 (ochre) from human spleen specimen. Dashed lines delineate the white pulp (WP). RP: Red pulp. Scale bar: 100 μm. (**G**) Scatter plot showing X and Y coordinates of segmented cells with MPO^high^CD117^high^ (red), MPO^high^CD117^high/med^ (pink), MPO^high^CD117^med^ (orange), MPO^med^CD117^low^ (green), MPO^low^CD117^low^ (blue, left panel). Ripley’s K function analysis of defined cell clusters (right panel).

Stress-induced myelopoiesis within the bone marrow alters progenitor spatial positioning with enhanced clustering (Herault et al., 2017) and generation of lineage-specific production sites (Wu et al., 2024; Zhang et al., 2021). Based on these studies, we profiled the spatial distribution of myeloid progenitors within the spleen and bone marrow (**Figure 3C-G**). We assessed the myeloperoxidase (MPO) expression pattern which represented a *bona fide* marker for differentiation from GMP towards mature neutrophils as found with flow cytometry analysis and intracellular staining (**Extended Figure 4D/E**) and immunofluorescence on sorted cells (**Extended Figure 4F**). EMH in the spleen was visible as isolated clusters within the red pulp, a structure not found in homeostasis (**Figure 3C, Extended Figure 4G/H**). Interestingly, neutrophils with an immature phenotype (dispersed MPO signal and reduced Ly6G expression, cluster 3) were situated close to their progenitors but distant to mature cells as compared to randomly distributed cells (**Figure 3D**), demonstrating the formation of neutrophil production sites around clustered myeloid progenitors. Furthermore, distance analysis between mature and immature neutrophils showed an increased spacing in the spleen as compared to bone marrow under control or CPM/AMD treatment (**Figure 3E**), thus confirming the disconnection between resident mature splenic neutrophils and the immature counterparts. To extend these analyses to human biology, we analyzed the proportion of myeloid progenitors in the human spleen by flow cytometry (**Extended Figure 5A**). Here, GMPs were present in the human spleen in a similar proportion as compared to human bone marrow and higher than in cord blood. Interestingly, human splenic GMPs differentiated preferentially into neutrophil-like cells as compared to monocytes (**Extended Figure 5B**), supporting the neutrophilic bias of splenic myeloid progenitors found in mice (**Extended Figure 2C**). To investigate the spatial distribution of myeloid progenitors and their descendants, we performed multiplexed imaging on spleens obtained from human organ donors (**Figure 3F/G, Extended Figure 5C-E**). Using histocytometry analysis, we clustered neutrophilic cells (CD11b^+^CD15^+^) into 5 groups based on the expression of MPO and CD117 (**Figure 3G**, **Extended Figure 5C-E**). We then used a Ripley’s K-function to assess whether the identified cells were clustered or randomly distributed (**Figure 3G**). Here, clusters with a progenitor (MPO^high^CD117^high^) or immature phenotype (MPO^high^CD117^high/med^, MPO^high^CD117^med^) displayed values that deviated from a homogenous Poisson distribution (Theoretical Poisson) supporting their clustered distribution (**Figure 3G**). Interestingly, these cell clusters are located in the red pulp (**Figure 3F**) and in close proximity to vessels (**Extended Figure 5D**).

### Epigenetic rewiring drives splenic neutrophil differentiation routes

To understand splenic differentiation dynamics, we performed trajectory inference analysis on our scRNA-seq. PAGA analysis revealed a main production and differentiation trajectory (Milestone 9 to 10, **Figure 4A** and **Extended Figure 6A-C**) but also a pronounced accumulation at early stages of the trajectory (**Figure 4B**) defined by a second path (Milestone 3, **Figure 4A**). Velocity analysis confirmed this differentiation pathway that deviated from the bone marrow main route (**Figure 4C/D**). Next, we questioned whether splenic neutrophils with an immature profile constituted an end-stage cell within the differentiation pathway. We hence sorted cells at the immature stage, and adoptively transferred them into a mouse under steady-state condition (**Figure 4E**). 16 hours after transfer, we assessed the expression of markers of maturation such as CD101 and CXCR2 (**Figure 4E**, **Extended Figure 6D**). Interestingly, we found that immature neutrophils originating from the bone marrow acquired expression of these markers, while splenic immature neutrophils failed to acquire maturation markers. In agreement with this intrinsic inability to adopt a mature phenotype, *in vitro* assessment of differentiation of immature neutrophils from bone marrow and spleen showed that spleen-derived cells had a reduced ability to fully mature of spleen-derived cells (**Extended Figure 6E**). Thus, these results culminate in a concept that neutrophils produced in the spleen rapidly differentiate into an end-stage immature subpopulation that constitutes the end of their maturation path.

**Figure 4.**
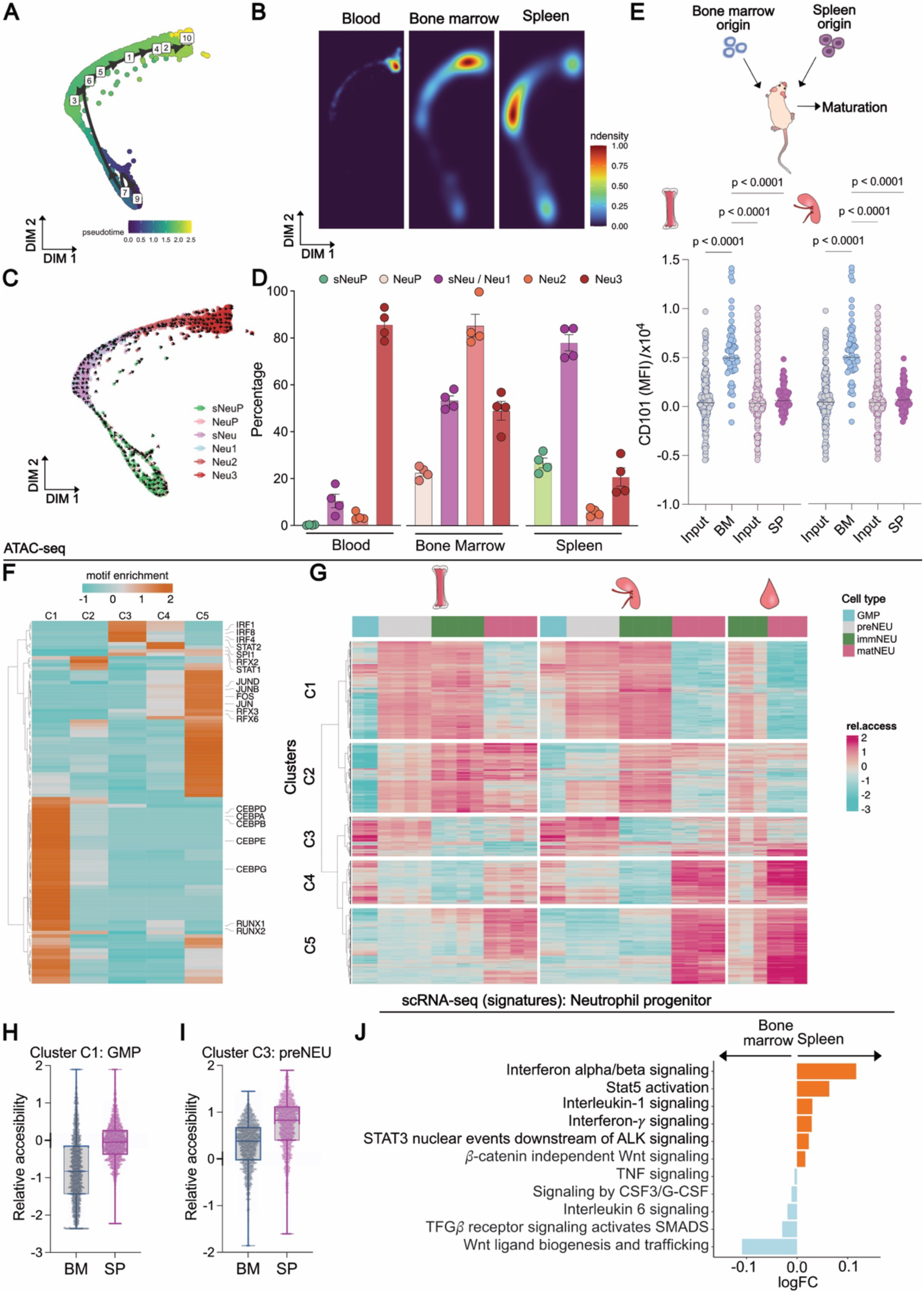
**Epigenetic rewiring of splenic neutrophil progenitors to generate alternative granulopoiesis differentiation paths**. C57BL6/J female mice were administered with cyclophosphamide (CPM), followed by daily administration of 5 mg/kg of AMD3100 (AMD). **(A)** PAGA trajectory plot calculated on aggregated neutrophils from blood, bone marrow and spleen representing pseudotime values. Milestones represent the dynamic transition between cell states. **(B)** Density plot showing cell density across neutrophil granulopoiesis obtained from scRNA-seq analysis in blood, bone marrow and spleen from CPM/AMD-treated mice. **(C)** Velocity analysis on scRNA-seq data from aggregated neutrophils from blood, bone marrow and spleen. **(D)** Relative abundance analysis of identified clusters of indicated organs. **(E)** Immature neutrophils (B220^-^CD19^-^CD3^-^CD115^-^F4/80^-^SSC^high^CXCR2^low^) isolated from bone marrow (dsRED) or spleen (Lyz2^eGFP^) were adoptively transferred intravenously into control mice. Flow cytometry analysis of CD101 expression of donor neutrophils isolated from the recipient bone marrow or spleen. Data is represented respective to CD101 levels of input. n = 20-200 cells out of 6 mice. One-way ANOVA analysis with Tukey’s correction. ****p<0.0001. **(F/G)** ATAC-seq analysis of sorted GMPs, preNEU, immNEU and matNEU from CPM/AMD-treated mice and indicated organs. (F) Heatmap showing motif enrichment analysis from unbiased clustered chromatin peaks. (G) Heatmap showing chromatin accessibility in indicated organs and cell types. **(H/I)** Analysis of relative accessibility peaks for cluster 1 in GMPs (H) or cluster 3 in preNEU (I). **(J)** Signature enrichment analysis in neutrophil progenitor cells from spleen and bone marrow for signaling pathways. Data are mean +/-SEM. GMP: Granulocyte-Macrophage Progenitor; immNEU: immature neutrophil; matNEU: mature neutrophil.

To obtain further insight into the mechanisms driving this accelerated differentiation, we profiled the epigenetic changes occurring at the stage of the neutrophil progenitor. We performed ATAC-seq (Assay for transposase-accessible chromatin followed by sequencing) in sorted GMPs, preNEU, immNEU and matNEU from the bone marrow, spleen, and blood (**Figure 4F/G, Extended Figure 6F/G**). Principal Component Analysis (PCA) showed that samples primarily clustered based on cell type (**Extended Figure 6F**). Subsequent unbiased clustering resulted in 5 main clusters of open chromatin regions (**Figure 4F/G**). Enrichment score analysis on these 5 clusters revealed an increased presence of motifs for CEBP transcription factor (TF) family (CEBPα, CEBPβ, CEBPδ, CEBPε, CEBPγ) and RUNX1 and RUNX2 in cluster 1 (C1) which is associated with neutrophil maturation (**Figure 4F**) (Khoyratty et al., 2021). Peaks of cluster 3 (C3) were enriched in motifs for Interferon Regulatory Factor (IRF, IRF1, 4 and 8) and STAT families, PU1 motifs for cluster 4 (C4) and AP-1 family motifs for cluster 5 (C5, JunD, JunB, Fos). As found by scRNA-seq, the chromatin accessibility of the maturation cluster C1 reflected the similarity between mature cells in the spleen and blood, suggesting their similar origin, and their striking difference with the immature splenic cells (**Figure 4G**). At the progenitor level, paired evaluation of general open peaks analysis showed an increased number of differential open chromatin regions in splenic GMPs, suggesting an active status at this level of development (**Extended Figure 6G**). These open peaks were particularly enriched in C1 cluster, supporting an active differentiation phenotype (**Figure 4H**). Of note, while no differences were found in the expression of the receptor for G-CSF, splenic GMPs showed increased levels of GM-CSF and IL-3 receptors previously associated with proliferative progenitors in emergency granulopoiesis (**Extended Figure 6H**) (Regan-Komito et al., 2020; Weber et al., 2015). On the other hand, C3 chromatin regions enriched for IRF and STAT TF families were more open in splenic preNEUs as compared to their bone marrow counterparts which accounted for an increased interferon signature in sNeuP (**Figure 4I**). To confirm this data with RNA levels, we performed enrichment score analysis on neutrophil progenitors in our scRNA-seq data (**Figure 4J**). Interestingly, pathway enrichment analysis showed that type I interferon and STAT signaling were particularly enriched in the splenic progenitors while TGFβ was reduced, which is in accordance with an enhanced progenitor differentiation and reduced regeneration and entrance into quiescence (Herault et al., 2017). Altogether, we find an interferon signature epigenetically imprinted in sNeuP which is associated with shortened differentiation paths to rapidly generate mobilizable end-stage neutrophils under emergency granulopoiesis.

### Splenic neutrophils propel host defense through interferon priming

We next sought to assess the functionality of spleen-derived neutrophils by profiling relevant neutrophil antimicrobial effector functions such as reactive oxygen species (ROS) production or Neutrophil Extracellular Trap (NET) formation. At transcriptomic level, splenic immature neutrophils showed enrichment in genes associated with NETosis, neutrophil activation, and ROS production (**Figure 5A, Extended Figure 7A/B)**. Furthermore, splenic neutrophils are characterized by increased expression of genes related to anti-bacterial or anti-fungal response. Our functional analysis confirmed these transcriptomic differences showing an increased ROS production (**Figure 5B**) in immature as compared to mature neutrophils within the blood compartment. In line with the increased ROS production, NETosis in response to LPS was augmented in circulating immature as compared mature neutrophils (**Figure 5C**). Of note, phenotypic analysis of FACS-sorted neutrophils showed that circulating immature neutrophils exhibited elevated expression of MPO and CD11b as compared to mature neutrophils (**Extended Figure 7C**). As expected, sorted neutrophils showed a prototypical nuclear band-shape for immature neutrophils irrespective of the organ of origin (**Extended Figure 7C**). On the other hand, granule protein content exemplified by myeloperoxidase (MPO, primary granules) or CD11b (tertiary granules) as well as cytoplasmic S100A8 showed no differences between bone marrow and splenic neutrophils. Contrary to observations made by others where immature neutrophils exhibit reduced effector capacity (Silvestre-Roig et al., 2019), we here show that splenic immature neutrophils produced during hematopoietic stress display similar or greater antimicrobial effector functions compared to neutrophils originating from the bone marrow despite their immature phenotype. We hence tested the antimicrobial abilities of sNeu in a model of UPEC-induced bladder infection (**Figure 5D**). Here, we analyzed the number of bacterial colonies in the bladder of CPM/AMD-treated mice in absence of the spleen and after adoptive transfer of sNeu. In line with their enhanced effector properties, lack of sNeu in splenectomized mice increased bacterial burden, an effect fully reverted by adoptive transfer of sNeu (**Figure 5E**). No changes were found in neutrophil numbers within the bone marrow, circulating or bladder (**Extended Figure 8A-C**).

**Figure 5.**
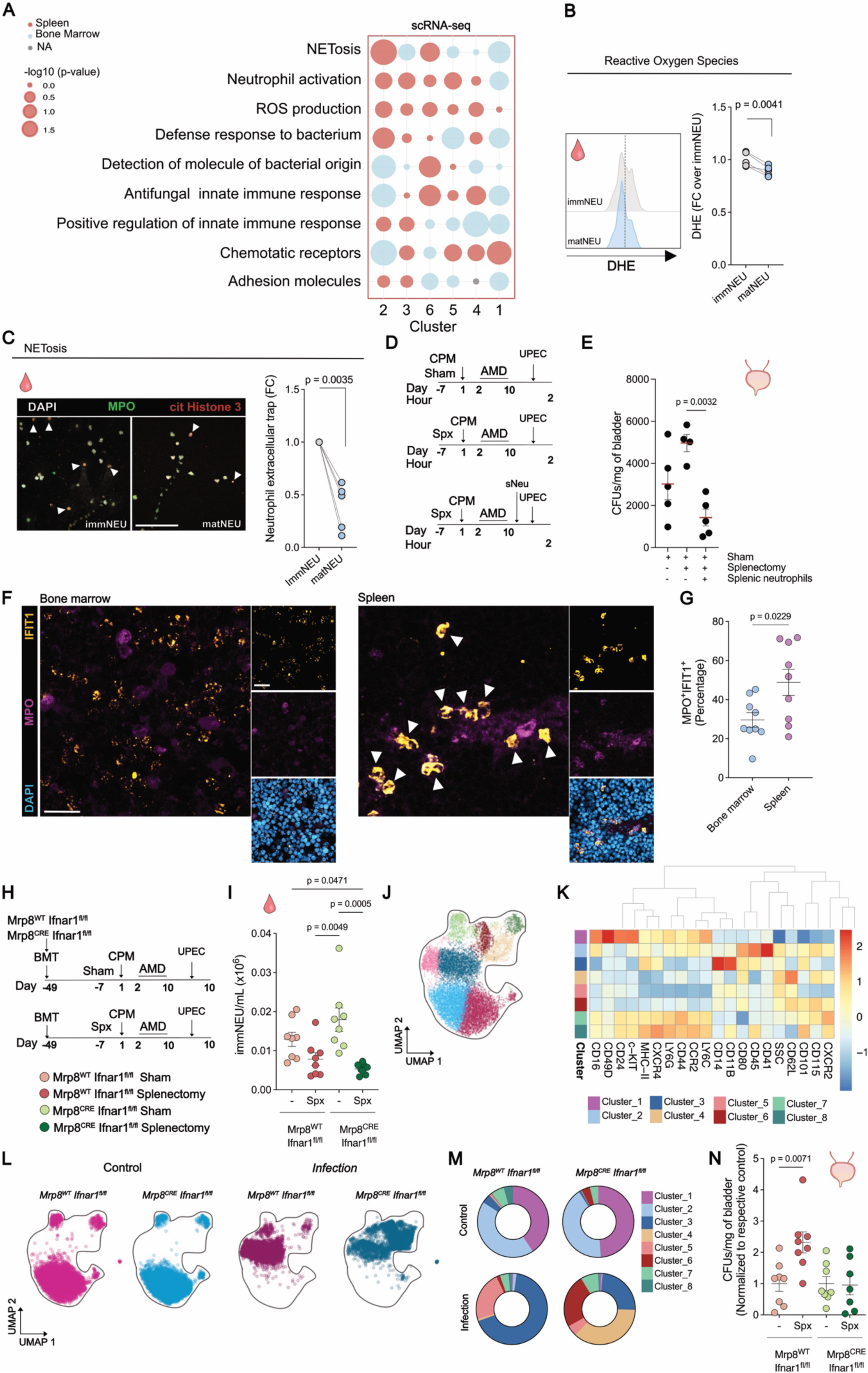
Interferon-primed splenic neutrophils facilitate bacterial clearance. C57BL6/J female mice were administered with cyclophosphamide (CPM), followed by daily administration of 5 mg/kg of AMD3100 (AMD). **(A)** Bubble plot showing the fold-change expression of indicated enriched signatures across clusters (2, 3, 6, 5, 4, 1 from immature to mature) between spleen (red) and bone marrow (blue). **(B)** Cytometry analysis of ROS using DHE in immature (Ly6G^+^CD101^low^CXCR2^low^) or mature (Ly6G^+^CD101^high^CXCR2^high^) blood neutrophils incubated with LPS (1 ng/mL). Two-sided paired t-test. **(C)** Confocal immunofluorescence analysis of NETs (DAPI (nuclei), MPO, citrullinated histone H3 (cit Histone H3)) in circulating neutrophils incubated with LPS (1 ng/mL). Scale bar: 100μm. **(D, E)** Mice were subjected to splenectomy (Spx) or sham operated. A group of splenectomized mice was administered with splenic neutrophils. All groups were infected with 5×10^8^ uropathogenic *E.coli* and sacrifice 2 hours post-infection. (D) Scheme of experimental procedure. (E) Bacteria colonies (colony forming units, CFU) measured in tissue bladder dissociates. **(F)** Confocal immunofluorescence of bone marrow and spleen tissue of CPM/AMD-treated C57BL6/J mice showing nuclei (DAPI, blue), Myeloperoxidase (MPO, magenta) and Interferon Induced Protein with Tetratricopeptide Repeats 1 (IFIT1, ochre). Scale bar: 20μm. **(G)** Analysis of the percentage MPO^+^IFIT1^+^ in the bone marrow and spleen. Two-sided unpaired t-test. **(H-N)** Mice were lethally irradiated and transplanted with bone marrow cells from *Mrp8^cre^Ifnar1^fl/fl^*or *Mrp8^wt^Ifnar1^fl/fl^*. After reconstitution, mice were subjected to splenectomy or sham, followed by treatment with CPM/AMD. **(H)** Scheme of experimental procedure. **(I)** Flow cytometry analysis of immature neutrophils in blood before infection. **(J)** UMAP showing unbiased clustering of aggregated cells from blood neutrophils before infection. **(K)** Heatmap analysis of surface proteins in identified clusters. **(L)** UMAP showing neutrophils across indicated genotypes and conditions. **(M)** Venn diagram showing cluster proportions in indicated conditions. **(N)** Bacteria colonies (colony forming-units, CFU) measured in tissue bladder dissociates. One-way ANOVA with Tukey’s correction. Data are mean +/-SEM.

To mechanistically explain the antimicrobial characteristics of spleen-derived immature neutrophils, we hypothesized that the elevated interferon signaling at the progenitor level (**Figure 4**) might confer enhanced antibacterial responses. Furthermore, we speculated that this interferon priming was restricted to the spleen environment as was previously shown for dendritic cells (Schaupp et al., 2020). In agreement with this idea, the proportion of progenitors expressing high levels of the interferon-responsive protein IFIT1 is enhanced in the spleen as compared to the bone marrow (**Figure 5F/G**). To causally test the importance of interferon signaling, we employed a mouse model lacking Interferon-alpha/beta receptor signaling in neutrophils (*Mrp8^cre^Ifnar1^fl/fl^*). *Mrp8^cre^Ifnar1^fl/fl^* and *Mrp8^wt^Ifnar1^fl/fl^* were subjected to splenectomy or sham treatment, followed by CPM/AMD application prior to bacterial infection (**Figure 5H**). Importantly, deletion of Type I interferon signaling in neutrophils did not affect the number of circulating neutrophils with an immature phenotype which were dramatically reduced after splenectomy (**Figure 5I**). On the other hand, analysis of surface marker expression in circulating neutrophils showed no differences in neutrophil activation before infection but a marked reduction in activated neutrophils upon bacteria administration (Cluster 3, **Figure 5J-M**). No changes in the absolute number of circulating mature neutrophils, total neutrophils, or monocytes before (**Extended Figure 8D-F**) or circulating immature neutrophils, mature neutrophils, or monocytes after infection (**Extended Figure 8G-I**) were detected between the two genotypes. Similarly, no differences were found in tissue-infiltrated neutrophils within the bladder (**Extended Figure 8J-M**). Confirming our previous, results splenectomy gave rise to a defective containment of the bacteria. Mice lacking Type I interferon signaling in neutrophils had an impaired ability to contain and clear bacteria, an effect not further accentuated when removing the spleen (**Figure 5N**). Collectively, these results demonstrate a role of the spleen environment for the priming of neutrophils produced in the spleen. Specifically, our data point towards the importance of Type I interferon signaling to maintain host defense fitness during stress-induced granulopoiesis.

## Discussion

Host immune responses against long-lasting stressors are associated with a profound remodeling of the hematopoietic compartment that is biased towards myeloid differentiation. Granulopoiesis is particularly prioritized under multiple continuous sterile and infectious challenges and constitutes an important use of energy that challenges the capacity of the bone marrow (Yvan-Charvet and Ng, 2019). Under certain inflammatory scenarios, granulopoiesis can be re-routed to other tissues that host hematopoietic progenitors that generate neutrophils; these potentially support the bone marrow production and maintain the immune response. Although generally considered a side-effect of persistent inflammation, the role of extramedullary neutrophil production and its contribution to host defense is understudied. Here, we identify an alternative yet accelerated neutrophil production in the spleen generating terminally differentiated cells with enhanced antimicrobial capacity contributing to pathogen containment. First, mapping stress-induced splenic granulopoiesis across a variety of perturbations characterized by low chronic inflammation revealed a generalized bias towards the generation of neutrophils with an immature phenotype. Second, splenic neutrophil progenitors are activated to rapidly differentiate into neutrophils that continuously exit their niche, feeding the circulating pool. Third, type I interferon priming of splenic progenitors links to production of neutrophils with enhanced effector function.

Persistent inflammation in unresolved infections initiates an evolutionarily conserved program of emergency granulopoiesis to cope with the need for neutrophils required to fight invading pathogens (Manz and Boettcher, 2014). Features of this programmed response are conserved across a variety of persistent low-grade inflammatory conditions, including cardiometabolic stress, cancer, and chronic infection (Bayne et al., 2012; Coffelt et al., 2015; Evrard et al., 2018; Li et al., 2022; Ohtsu et al., 2000; Regan-Komito et al., 2020; Silvestre-Roig et al., 2020; Zhu et al., 2018). Among these features, the elevated production of immature neutrophils and their premature release to the circulation is a hallmark of these pathologies and is commonly associated with disease severity (Carissimo et al., 2020; Mackey et al., 2019; Prada-Medina et al., 2020; Schulte-Schrepping et al., 2020; Silvin et al., 2020; Wan et al., 2023). In addition to confirming the mild expansion of neutrophil progenitors and immature neutrophils within the bone marrow, we here identify a strikingly increased granulopoiesis within the spleen peaking at the stages of preNEU and immature neutrophils. The similar proportion of these subpopulations within the bone marrow also suggests that both granulopoiesis pipelines are governed by common differentiation hierarchies primarily arising at the level of the preNEU as previously suggested (Evrard et al., 2018). However, the rate of cell differentiation differs significantly between the two niches, with splenic neutrophil progenitors outperforming the capacity of their bone marrow counterparts. Our trajectory analysis also shows that this heightened progenitor activity is accompanied by a more rapid differentiation route to generate and release effector neutrophils in a shorter time frame. These two processes (cell differentiation and mobilization) might have evolved in parallel within the spleen environment to efficiently cope with neutrophil demands and to accelerate the elimination of the invading pathogens. On the other hand, the activation of this program under chronic sterile inflammatory perturbations is probably associated with the previously observed pathogenic contribution of spleen-derived myeloid cells in mice (Cortez-Retamozo et al., 2012; Dutta et al., 2015; Regan-Komito et al., 2020; Robbins et al., 2012) and humans (Emami et al., 2015; Kim et al., 2014). Moreover, our study also reveals the disparate origins of splenic neutrophils. The disruption in cell maturation during splenic granulopoiesis with immediate exit to the bloodstream indicates that the identified mature neutrophil population in the spleen is not produced on site. Our data demonstrate a phenotypic, spatial, and functional separation between immature and mature splenic subsets in both mouse models and in human spleen specimens. Therefore, we propose that this mature splenic population is in fact blood-borne, and probably contributes to the local function of these subpopulations or the clearance of circulating aged neutrophils. Consequently, future functional studies on neutrophils produced during EMH should carefully discriminate between discrete subsets when comparing neutrophils of different maturation states and origin.

Our observations support a stress-driven granulopoiesis even under situations of low chronic inflammation. While the existence of the immature neutrophils in the circulation (“left-shift”) is commonly associated with an increase of discharge-inducing signals that surpass the signaling threshold necessary for inducing neutrophil mobilization to the blood, their appearance under chronic stress cannot be explained by a compensatory mechanism to meet the demand of cells. In fact, the release of these cells might be a consequence of (1) a collateral effect of a dysfunctional granulopoiesis or (2) an active process to deal with the instigating stimulus. For the first possibility, our transcriptomic and epigenetic data shows that splenic GMPs have an increased number of open chromatin peaks at cluster 1, which is enriched for TFs involved in differentiation and maturation as compared to those found in the bone marrow, supporting an activated and differentiative phenotype as previously suggested (Coppin et al., 2018). This epigenetic and transcriptional status explains the observed elevated proliferation and differentiation capacity and might consequently result in the generation of neutrophils with reduced maturation. Furthermore - and despite stem cells and progenitors exhibiting similar levels of G-CSF receptor in both organs-, HSPCs of splenic origin displayed increased levels of IL-3 and GM-CSF receptors. This may elevate their responses to growth factors involved in stem cell survival and proliferation during emergency hematopoiesis (Regan-Komito et al., 2020; Weber et al., 2015) and thus support an enhanced intrinsic splenic neutrophil progenitor responsiveness and activity. Our spatial analysis also revealed that splenic neutrophils originate from clustered production sites in the red pulp, an active process that is part of the emergency granulopoiesis program, as previously suggested (Herault et al., 2017). Notably, this clustered production of neutrophils within the red pulp can also be inferred from our analyses of human spleen samples obtained from organ donors, suggesting that this process is conserved across species. However, the functional consequences of this process in human infectious diseases, cancer, or metabolic pathologies remains to be studied.

On a functional level, our results illustrate an alternative differentiation pathway in the spleen to rapidly generate fully competent neutrophils. In contrast to their immature phenotype, spleen-derived neutrophils are fully armed with antimicrobial effector functions that help to contain and eliminate invading bacteria. Contrasting data showing reduced (Drifte et al., 2013), similar (van Grinsven et al., 2019) or increased (Hsu et al., 2019; Rice et al., 2023) functional properties of phenotypically immature neutrophils suggest that the function of these cells may be context-and origin-dependent. Using adoptive transfer strategies, we here provide causal evidence for the beneficial role of spleen-borne immature neutrophils in the context of infections, yet genetic tools to modulate the levels and functions of this subpopulation are necessary to dissect their role in greater depth. Importantly, the pathogen-containing properties of spleen-derived immature neutrophils are in agreement with the concept of an active, on-demand production of this subpopulation in response to inflammatory stress. Mechanistically, we demonstrate that the spleen can host the production of these neutrophils due to its niche characteristics and the ability of progenitor priming through type I interferons. Given the functionality of these cells, it is plausible that the selective mobilization of neutrophils at different stages of maturation might help adapt the host neutrophil circulating pool to specific inflammatory responses depending on the functional needs. Such concept is exemplified by the program of “neutrophil aging”, which generates cell subpopulations that meet different functions across the day and in synchrony with the higher likelihood of exposure to external challenges during the activity phase (Adrover et al., 2020; Adrover et al., 2019). This functional plasticity can also be achieved by releasing neutrophils with various degrees of maturation exhibiting distinct epigenetic and metabolic properties, allowing them to adapt to different environments counterparts (Hsu et al., 2019), or to strongly react to invading pathogens under life-threatening conditions.

The coexistence of mature and immature neutrophil subsets in altering proportions under different pathological conditions may lead to different disease outcomes and is likely to be disease-dependent. On the other hand, in non-infectious diseases, the existence of this subpopulation could cause undesired effects due to its persistent action in the absence of an invading pathogen. However, this possibility remains to be investigated in future studies. In conclusion, we show that the spleen serves as an immune-supporting platform to produce neutrophils with enhanced antimicrobial function to cope with infections under prolonged stress. Our findings illustrate a host mechanism that exploits increased neutrophil plasticity and cell maturation as a source of functional heterogeneity to generate rapid and effective responses against invading pathogens.

## Acknowledgements

We thank Ulrich Dobrindt (University of Münster) for providing the uropathogenic *E*.coli strain used for the infection experiments. This project was supported by the Else Kröner Fresenius Stiftung (2017_A13) and the Deutsche Forschungsgemeinschaft (CRC TRR332 project A1). C.S.R. receives support from IZKF of University of Münster. O.S. receives support from the Deutsche Forschungsgemeinschaft (CRC TRR332 projects A2 and Z1, CRC1123 project A6, CRU342 project 1, project 502158695, SO876/16-1), the Leducq Foundation, Novo Nordisk, the EU (PRAETORIAN Doctoral Network), and the IZKF and the IMF of the University of Münster. D.R.E. is supported by the Deutsche Forschungsgemeinschaft (CRC TRR332 project A3). J.J. is supported by the Deutsche Forschungsgemeinschaft (CRC TRR332 project A5). T.C. is supported by the Deutsche Forschungsgemeinschaft (CRC TRR332 project B4).

## Author contributions

C.S.R., R.C., and O.S., conceptualized the study and designed experiments. C.S.R., performed most of the experiments and analyzed data. R.C., A.B., performed experiments and analyzed data. L.M.V., A.H., M.F., M.R. Q.B., M.G., J.S., S.S., P.L., C.O.S., performed experiments. M.R., helped with MACsima image acquisition and analysis. D.A., C.M., helped with mass cytometry (CyTOF) panel design and generation, and data acquisition. D.R.E., helped with intraurethral infection model. M.G., J.S., A.C., designed, performed and acquired data of flow cytometry analysis of human spleen and in vitro differentiation of splenic progenitors. A.K., provided human spleen specimens for imaging. J.J., provided scRNA-seq dataset for murine oropharyngeal carcinoma. C.S.R., and O.S., wrote the manuscript. R.C., A.B., M.R., P.D., T.C., edited and provided comments to the manuscript. C.S.R., and O.S., funded the project.

## Cell lines

The murine oropharyngeal carcinoma cell line MOPC (C57BL/6-derived, HPV16 E6/E7-) was obtained from Dr. William Chad Spanos and John H. Lee (Sanford Research/University of South Dakota, Sioux Falls, SD, US) (Hussain et al., 2022; Klein et al., 2017; Williams et al., 2009). The cells were cultivated in a special medium (67% DMEM, 22% Hams F12 nutrient mix, 10% Fetal Bovine Serum, 1% penicillin-streptomycin, 0.5 µg/ml Hydrocortisone, 8.4 ng/ml Cholera Toxin, 5µg/ml Transferrin, 5 µg/ml Insulin, 1.36 ng/ml Tri-Iodo-Thyronine, 5 µg/ml E.G.F.). During cultivation, the cell lines were regularly tested for mycoplasma contamination with negative results. B16-F10 cells were cultured in DMEM (Gibco/Life Technologies) supplemented with 10% Fetal Bovine Serum and 1% penicillin-streptomycin and incubated at 37 °C, 5% CO2. During the exponential phase of cell growth, after 2 days of culture, cells were detached by 0.5% trypsin and EDTA (Gibco/Life Technologies) and washed with PBS before injection. All cells were grown in a monolayer at 37 °C in a humidified incubator with 5% CO2.

## Ethics statement

All mouse experiments were performed according to European guidelines for the Care and Use of Laboratory Animals. Protocols were approved by the Committee on the Ethics of Animal Experiments of the Regierung von Oberbayern and LANUV.

Splenic biopsies from organ donors were collected from leftover material from the Institute for Transfusion Medicine at the Medical School Essen, as approved by the ethics committee (05-2768) of the Medical Faculty of the University of Duisburg-Essen, Germany.

The use of blood and tissues was approved by the Ethical Committee for Clinical Investigation of the Hospital del Mar Research Institute (CEIC-IMIM 2011/4494/I and 2014/5892/I). Fresh samples were obtained from the Mar Biobanc tissue repository with patient signed informed consent. All the blood and tissue samples were coded and relevant clinical information remained anonymous.

## Mouse procedures

For mouse experiments, statistical power calculations were performed as described at http://www.stat.uiowa.edu/~rlenth/Power/ to determine sample size. Mice were assigned to groups randomly and data collection and analysis were performed blinded. Mice were housed according to institutional regulations with ad libitum access to food and water. All mice used were males or females in the C57BL/6J background from 8-20 weeks-old. Aged mice were used at the age of 84-96 weeks-old. Apoliprotein E-deficient (*Apoe^-/-^*), Tg(CAG-DsRed*MST)1Nagy, Dendra2 (129S-Gt(ROSA)26Sortm1(CAG-COX8A/Dendra2)Dcc/J), *Ifnar1^-/-^*, B6.Cg-Tg(S100A8-cre,-EGFP)1Ilw/J (Mrp8^cre^), *LyzM^GFP^* were purchased from the Jackson Laboratory. Mrp8^cre^ was intercrossed with *Ifnar1^fl/fl^*mice to generate *Mrp8^cre^ Ifnar1^fl/fl^*.

### Mouse model of CPM/AMD

Female mice were injected intraperitoneally with 4 mg of cyclophosphamide followed by daily subcutaneous injection (10 to 21 days) of AMD3100 (5 mg/kg, Biorbyt).

### Mouse model of hypercholesterolemia

Female *Apoe^-/-^*mice were fed with high-cholesterol (21% fat and 0.15% cholesterol) diet for 16 weeks.

### Mouse model of melanoma cancer

Female mice were injected subcutaneously in the left flank with 50000 B16 tumor cells in 30μL of Matrigel (Gibco). Mice were sacrificed 4 weeks after tumor cell administration.

### Mouse model of murine oropharyngeal carcinoma

The MOPC cells were injected subcutaneously (s.c. 1 x 10^6^ in 100 µl PBS) into the flank of C57BL/6 as described previously (Pylaeva et al., 2022).

### Mouse model of chronic endotoxemia

Female mice were injected intraperitoneally with 10 µg of LPS (*E.coli* K12, Invivogen) for 3 consecutive days.

### Splenectomy

Mice were anaesthetized with Midazolam (5,0 mg/kg), Medetomidin (0.5 mg/kg) and Fentanyl (0.05 mg/kg). After exposure of the spleen on the left flank with a 1 cm cut, the splenic artery and vein are cauterized, and the spleen is removed. Then, the peritoneal tissue and the skin were closed using 6.0 suture. The anaesthesia was antagonized by subcutaneous administration of Atipamezol (2.5 mg/kg) and Flumazenil (0.5 mg/kg).

### Bromodeoxiuridine (BrdU) pulse assay

For BrdU assays, mice were injected intraperitoneally with 1 mg of BrdU and sacrificed at indicated time-points. BrdU incorporation was measured on isolated cells by flow cytometry using the BrdU flow cytometry kit (BD Pharmigen) following manufacturer’s instructions.

### Partial irradiation

Anaesthetized mice were placed in the irradiator and the abdominal area was covered with a 1.375” diameter shield (Braintree Scientific) and irradiated with 1x 6.5 Gy.

### Bone marrow transplant

For bone marrow transplant, mice are lethally irradiated with 1x 7.5 Gy followed by reconstitution with 2×10^6^ bone marrow cells. Bone marrow reconstitution is followed 6 weeks after irradiation.

### Intrasplenic and intratibial injection of cells

Mice were treated with 4 mg of cyclophosphamide to favor stem cell engraftment. After 24 hours, mice are anaesthetized with Midazolam (5.0 mg/kg), Medetomidin (0.5 mg/kg) and Fentanyl (0.05 mg/kg). For intrasplenic injection of stem cells, after the exposure of the spleen on the left flank with a 1 cm cut, 4×10^3^ lineage negative cells were injected subcapsulary. For intratibial injection, the bone was perforated with a XXG needle and 1×10^5^ lineage negative cells were slowly injected. Lineage negative cells were obtained from the bone marrow cells subjected to Lineage depletion kit (Miltenyi).

### *In vivo* maturation assay

Immature neutrophils (B220^-^CD19^-^CD3^-^CD115^-^F4/80^-^ SSC^high^CXCR2^low^) isolated from bone marrow (dsRED) or spleen (Lyz2-eGFP) were adoptively transferred intravenously into control mice. Flow cytometry analysis of CD101 expression of donor neutrophils isolated from the recipient bone marrow or spleen.

### Intraurethral infection

Mice are anaesthetized with Midazolam (5.0 mg/kg), Medetomidin (0.5 mg/kg) and Fentanyl (0.05 mg/kg). 5×10^8^ uropathogenic *E.coli 536* were inoculated intraurethrally and colony forming unit were measured in CPS Elite agar plates (Biomerieux) 2 or 4 hours after infection.

### Human cell processing

Peripheral blood mononuclear cells (PBMCs) were isolated from whole blood collected with EDTA anticoagulant via Ficoll–Paque Premium (Cytiva,) following the manufacturer’s instructions. PBMCs were counted using Turk solution, resuspended in fetal bovine serum (FBS; Gibco) with 10% dimethyl sulphoxide (DMSO, Sigma-Aldrich). Umbilical cord blood (CB) was obtained from healthy full-term newborns in 10 ml heparinized tubes (BD Vacutainer) and cells were isolated via Ficoll–Paque Premium.

Human splenocytes were obtained from fresh tissue samples as reported in published studies (Magri et al., 2014). Briefly, mononuclear cells were isolated from histologically normal spleens from deceased organ donors or trauma patients without clinical signs of infection or inflammation that had undergone splenectomy.

Bone marrow cells were obtained from total hip arthroplasty surgeries after cutting coronally in 4-6 mm thick slices. 1 mm of minced tissue was digested in filtered 4 mg/mL dispase II (Sigma), and 2 mg/mL collagenase I (Worthington Biochem) in PBS for one hour on a rocker at 37 °C. The cell suspension was then filtered through a 100 µm cell strainer (Fisher) and remaining tissue was washed five times in culture medium (AlphaMEM with 15 % FBS, 1% P/S, 0.1 mM LAA, 1 mM L-glutamine) and strained. For human cell suspensions, cells were treated with red blood cell lysis buffer (eBioscience) before flow cytometry staining or culture.

### Colony-Forming Unit assay

Human progenitors were isolated from frozen splenocytes collected after collagenase treatment (Magri et al., 2014). Splenic progenitor populations (hematopoietic stem cell; lineage^-^CD45^med^CD34^+^), MEP (myeloid-erythroid progenitor; lineage^-^ CD45^med^CD34^+^CD38^+^CD45R^-^IL3Ra^-^), CMP (common myeloid progenitor; lineage^-^ CD45^med^CD34^+^CD38^+^CD45R^-^IL3Ra^+^), GMP (granulocyte-macrophage progenitor; lineage^-^CD45^med^CD34^+^CD38^+^CD45R^+^IL3Ra^+^) were sorted on a fluorescence-activated cell sorter FACSAria (BD Biosciences) using FACSDiva software (BD Biosciences). Cell-sorted cells were plated in MethoCult^TM^ Optimum (H4035, STEMCELLS Technologies) supplemented with IL-3, GM-CSF, G-CSF and SCF. Plates were incubated at 37 °C for 14 days. Afterwards, Colony-Forming Units (CFC) were collected from the plates by flushing the agar in RPMI medium. Cells were washed twice by centrifugation and then stained with fluorochrome-labelled antibodies.

### Mouse tissue processing

Mice were euthanized by ketamine/xylazine overdose, the blood was collected by heart puncture after which the mice were flushed with 10 ml of ice-cold PBS. Subsequently, organs were harvested and directly processed for functional assays or flow cytometry analysis. Blood was collected in a tube containing EDTA and incubated with erythrocyte lysis buffer (Biolegend) for 6 minutes on ice for erythrocyte lysis. After washing, cells were centrifugated and resuspended in HANKs buffer (HBSS w/o Mg and Ca + 0.06% BSA + 0.3mM EDTA). For isolation of bone marrow cells, the femur was harvested and flushed using 5 mL of HANKs buffer using a syringe with a 21G needle. Splenic cells were obtained by the mince and smashing of the spleen over a 70um filter. Bone marrow or splenic cells were subjected to erythrocyte lysis in 1mL of erythrocyte lysis buffer (Biolegend) for 1 minute on ice. After washing, cells were centrifugated and resuspended in HANKs buffer. For lung and bladder tissue processing, weighted tissue was minced and digested in 1.25mg/mL Liberase TM + 1U/mL Dnase I for 30 minutes at 37°C. After filtering through a 30uM filter, tissue dissociated cells were incubated in RPMI +1%FCS for 30 minutes. After washing in HANKS buffer, cells were subjected to erythrocyte lysis in 1mL of erythrocyte lysis buffer (Biolegend) for 1 minute on ice. After washing, cells were centrifugated and resuspended in HANKs buffer.

### Cytospin

FACS-sorted mature (B220^-^CD19^-^CD3^-^CD115^-^F4/80^-^SSC^high^CXCR2^high^) or immature (B220^-^CD19^-^CD3^-^CD115^-^F4/80^-^SSC^high^CXCR2^low^) from blood, bone marrow or spleen were centrifugated at 270g for 5 minutes using the centrifuge Cellspin III (Thermac). After fixation with PFA 4%, cells were processed for immunohistochemistry as described below.

### Immunohistochemistry

For immunohistochemistry analysis, tissue was fixed in 3 mL PLP overnight at 4°C. After washing 2x times in phosphate buffer, tissue was dehydrated by successive sucrose gradients (10%, 20%, 30%) for 2 hours at 4°Cand then embedded in Tissue Tek O.C.T. compound (Sakura Finetek) for analysis. 7μm sections were then prepared for immunohistochemistry analysis. Sections were rehydrated in Tris 0,1M (pH 7.4), followed by a permeabilization step in Tris 0,1M (pH 7.4) + 1% Triton X100 for 1 hour at RT. After blocking for 1 hour at RT in blocking solution (Tris 0,1M (pH 7.4) + 1% BSA + 0,3% Triton), sections were incubated with primary antibodies at indicated dilutions (see antibody list) in blocking solution o.n. to 72 hours at 4°C in the dark. After washing 2x in Tris 0,1M (pH 7.4), sections were incubated with secondary antibodies for 2 hours at RT in blocking solution. Finally, and after washing, sections were stained with 4′-6′-diamidine-2′-phenylindole (DAPI, 1:1000, Invitrogen) and mounted in Prolong (Thermofisher) before imaging.

### MACSima Imaging Cyclic Staining (MICS)

Multiplex immunohistochemistry of spleens from human donors was performed using a MACSima imaging system (Miltenyi Biotec). In brief, cyclic immunofluorescence imaging consisting of repetitive cycles of staining using immunofluorescent antibodies, imaging of different regions, and erasure of fluorescence by photobleaching was performed.

Cryosectioned, PFA-fixed human spleens mounted on microscopy slides were blocked using a blocking buffer containing 10% BSA and 2% goat serum for 1h at RT in MACSwell sample carriers before nuclei were counterstained with DAPI. Samples were then placed into the MACSima imaging system, where they underwent repetitive cycles of staining using immunofluorescent antibodies, multi-field fluorescent imaging, and signal erasure by photobleaching. Neutrophils were identified as CD15+ (VIMC6, Miltenyi Biotec, FITC, 1:50) CD11b+ (REA1321, Miltenyi Biotec, PE, 1:50) cells and their phenotype and maturation status characterized using antibodies against MPO (REA491, Miltenyi Biotec, FITC, 1:50) and CD117 (REA787, Miltenyi Biotec, PE, 1:50). The tissue environment was characterized using antibodies against Vimentin (REA409, Miltenyi Biotec, PE, 1:50), CD45 (5B1, Miltenyi Biotec, APC, 1:50) and CD34 (AC136, Miltenyi Biotec, PE, 1:50). Nuclei were counterstained with DAPI before samples were placed in the MACSima imaging system.

## Data analysis and visualization

MACsima images were stitched and preprocessed using MACS iQ View Analysis Software (Miltenyi Biotec). For downstream analysis of confocal and MACsima images, images were subjected to Gaussian Blur filtering (factor = 4-6), then cells were segmented based on the DAPI signal using the StarDist plugin (Schmidt et al., 2018) (Probability score = 0.3; Overlap score = 0.2) in ImageJ (NIH) followed by the donut algorithm (distance = 12 pixels) in MACS iQ View. For confocal images in Figure 3C, segmented cells were gated using the scatter plot analysis and plotting average intensity of myeloperixodase (MPO) and Ly6G signal: myeloid progenitors (MPO^high^ y6G^low^), immature (MPO^med^Ly6G^low-med^) and mature (MPO^low^Ly6G^high^) neutrophils. Distance analysis was measured using the distance analysis function in the MACS iQ View Analysis Software. For confocal images in Figure 5G/H, segmented cells were gated based on MPO and IFIT expression using the scatter plot analysis function, and the number of double positive cells was analyzed. For human spleen samples, segmented cells were gated using the scatter plot analysis and gated on average cell intensity of CD11b and CD15. CD11b+CD15+ were then further gated using the scatter plot analysis of MPO and CD117 average cell intensities. Five populations were obtained: MPO^high^CD117^high^, MPO^high^CD117^high/med^, MPO^high^CD117^med^, MPO^med^CD117^low^ and MPO^low^CD117^low^. Gated populations were exported as fcs and loaded using the Flowcore package in R. Ripley’s K function analysis was then calculated using the *spatsat* package and plotted using plot() function from package *stats4* to assess the clustering of populations of interest.

### Flow cytometry

Mouse cells were stained with fluorescent-labelled antibodies in staining buffer (Biolegend, 20 min, 4 °C). Cells were fixed and permeabilized for intracellular cytokine staining using the Cytofix/Perm buffer (Bioleged). Flow cytometry was performed using the LSR Fortessa (Beckton Dickinson) or Cytek Aurora (Cytek) and data was analysed using FlowJo software (Beckton Dickinson). For human cells, cells were incubated with Fc blocking reagent (Human TruStain FcX, BioLegend) and appropriate labeled mAbs (**Table 1**) for 20 min at 4°C. Dead cells were excluded with DAPI. Cells were acquired with an LSR Fortessa or FACSCalibur flow cytometers (BD Biosciences) and data analyzed with FlowJo software (TreeStar).

### scRNA-seq

Single cells suspensions from blood, bone marrow and spleen were analyzed with flow cytometry to obtain total cell counts per sample. Samples were then normalized across conditions to obtain similar number of target cells (myeloid cell populations and progenitors). Normalized samples were then incubated with hashtag antibodies (see **Table 1**) specific for each organ, mouse and condition. After extensive washing, cells were pooled FACS-sorted using the following gating strategy: DAPI^-^ CD115^+^ Ly6C^+^ and enriched with cKit^+^ cells as described in Extended Figure 2A. Sorted cells were then loaded into the 10x Genomic platform and processed with 10X Chromium Next GEM Single-Cell 3′ Reagent Kit with Feature Barcoding technology following manufacturer’s instructions. Libraries were sequenced at the IMGM sequencing platform (Munich) using a lllumina NovaSeq S2 Run (128 cycles). Raw reads were processed with cellranger 3.1 and aligned with the referenced transcriptome Mus Musculus GRCh38. Control and CPM/AMD cells were loaded in two independent days, and obtained libraries were processed at the same time.

Data processing was performed with R and SingleCellExperiment. Raw counts were filtered using isOutlier function from scater package for library size (log-transformed library sizes), log-transformed number of genes per cell, and mitochondrial content to identify cells with values below 3 times the median absolute deviation. Low-abundance genes and genes expressed by less than 5 cells were excluded. Hashtag identification and removal of doublets and multiplets were performed using CiteFuse. Batch correction across experimental days was performed using fastMNN and bachelor package.

Counts were log normalized using lognormcounts() function. For dimensionality reduction, we used the Uniform Manifold Approximation and Projection (UMAP) method as implemented in the runUMAP (n = 10) function from the Singlecellexperiment package. Specifically, we reduced the dimensions of the dataset to 10 using PCA analysis on highly variable genes (calculated on top 20% genes), resulting in a lower-dimensional embedding. This step was performed prior to correcting for batch effects within the dataset. To account for batch variation, we employed the buildSNNGraph function with default parameters, which constructs a shared nearest neighbor graph based on the Jaccard similarity index between cells. The final clusters were then identified through the application of the Louvain community detection algorithm via the cluster_louvain function from the igraph package.

Cell annotation was performed using *cellassign* package and curated analysis based on previously published datasets defining neutrophil subpopulations (Evrard et al., 2018; Kwok et al., 2020; Muench et al., 2020).

## Signature calculation

To obtain the signature from immature splenic or bone marrow neutrophils, condition labels were assigned to facilitate pairwise comparisons. Cells were distinguished based on their organ of origin, blood, bone marrow, and spleen and treatment type, vehice or CPM/AMD. Conditions were defined for each combination of cell type, organ, and treatment. aggregateAcrossCells() was employed to aggregate data across individual cells by mouse and condition. Genes expressed in at least 10% of the cells in each condition were retained. For differential expression analysis, the gene-filtered dataset was input into an edgeR DGEList object for normalization. Cells with fewer than 10 observed events were discarded and normalized using *calcNormFactors().* Then, a design matrix capturing condition effects was constructed, and dispersion was estimated using estimateDisp() to account for biological variability. The glmQLFit() function was used for a general linear model fitting. A series of contrasts were defined to test generalized linear model quasi-likelihood F-tests (glmQLFTest() function).

### Signature enrichment pathway analysis

Signature enrichment calculation was performed in group of 1000 cells, using Ucell function (Andreatta and Carmona, 2021) integrated on escape package (Borcherding et al., 2021). The following gene lists were used for signature calculation: Interferon alpha/beta signaling (R-HSA-909733), STAT5 activation (R-HSA-9645135), interleukin 1 signalling (R-HSA-9020702), interferon gamma signalling (R-HSA-877300), STAT3 nuclear events downstream the ALK signalling ( R-HSA-9701898), beta catenin-independent Wnt signalling (R-HSA-3858494), TNF signalling (R-HSA-75893), signalling by CSF3-G-CSF (R-HSA-9674555), IL-6 signalling (R-HSA-1059683), TGF-beta receptor signalling activates SMAD (R-HSA-2173789), Wnt ligand biogenesis and trafficking (R-HSA-3238698), defense response to bacterium (GO:0042742), detection of molecule of bacterial origin (GO:0032490), positive regulation of innate immune response (GO:0045089), and antifungal innate immune response (GO:0061760). Other gene lists are described in **Supplementary_Table_1**.

### Trajectory and velocity analysis

PAGA tree analysis was performed on aggregated blood, bone marrow and splenic cells from CPM/AMD group using the function ti_paga_tree() from the Dynverse package collection (Saelens et al., 2019). https://github.com/dynverse/dyno. Velocity calculation was performed using scVelo (Bergen et al., 2020) and Velocyraptor packages.

### Mass cytometry

All directly conjugated antibodies were purchased from Standard Biotools. Conjugation of additional purified unlabeled antibodies was performed using the Maxpar X8 Metal Labelling Kit (Standard Biotools) according to manufacturer’s instructions. All antibodies were titrated to determine the optimal staining concentration.

Blood, spleen, and bone marrow cells were stained with the panel of antibodies outlined in **Table 1**. First, one million cells per sample were stained with rhodium DNA intercalator (Cell-ID™ Intercalator-Rh, Standard Biotools) to distinguish live/dead and barcoded using the Cell-ID 20-Plex Pd Barcoding Kit (Standard Biotools) as per manufacturer’s instructions. Samples from each tissue source were combined to create one multiplexed sample and incubated with Fc receptor blocking reagent prior to staining with a cocktail of cell surface antibodies for 30 minutes at room temperature. Stained cells were washed once in Cell Staining Buffer (CSB), fixed for 10 minutes in 1.6% PFA (Thermofisher), and incubated overnight with an Iridium DNA intercalator (Standard Biotools).

Stained samples were washed with CSB and cryopreserved in 1ml FCS containing 10% DMSO. Samples were stored at-80 until acquisition. On the day of acquisition, samples were thawed, washed twice in CSB supplemented with benzonase and washed twice in Maxpar Cell Acquisition Solution (Standard Biotools). For acquisition, samples were resuspended in CAS containing EQ 4 Element Calibration Beads (Standard Biotools) and acquired at a rate of < 350 events/second on a Helios Mass Cytometer. After acquisition, data (.fcs files) was normalized, concatenated, and debarcoded using the instrument’s CyTOF Software v7.0.

Analysis was performed using Flowjo software and FlowCore and Cytometry dATa anALYsis Tools (CATALYST) packages. Using Flowjo software, samples were gated for live cells and downsampled. Processed fcs files were loaded using FlowCore package and transformed using arcsinh transformation with a cofactor of 5 using the asinh() function. The working object was generated using prepData() function. For visualization, tSNE plots were generated using runTSNE() on a subset of 5000 cells. Unsupervised clustering was generating using FlowSOM and ConsensusClusterPlus packages. Cell annotation was performed based on main lineage-defining surface markers (B cells: B220; T cells: CD3^+^; NK cells: NK1.1^+^; Neutrophils: CD11b^+^Ly6G^+^; Macrophages: CD11b^+^F4/80^+^; Monocytes: CD11b^+^, CD115^+^; Stem cells: cKit^+^).

### ATAC-seq

ATAC-seq libraries were prepared from ex vivo sorted cells (7.000-50.000 cells) following the OmniATAC protocol (Corces et al., 2017) but using KAPA HiFi HotStart ReadyMix (Roche) for PCR amplification. PCR products were cleaned up twice with AMPure XP beads (Beckman Coulter) at a ratio of 1.2x and eluted into 20µl. Libraries were sequenced on Novaseq 6000 at 2×150bp.

Reads were trimmed from Nextera adapters using fastp, then mapped to GRCm38 using bowtie2 (-X 2000--very-sensitive), duplicate-marked with samblaster, and filtered for primary alignments to the main (chr1-19,X,Y) chromosomes, non-duplicates and a MAPQ > 20.

Peaks were called per sample with macs2 (--keep-dup=all--nomodel--extsize 100--shift-50--min-length 250) and filtered against a combined blacklist of common NGS artifacts (Amemiya et al., 2019) available at https://github.com/Boyle-Lab/Blacklist/tree/master/listsformm10) and mitochondrial gene homologs in the mouse genome ((Buenrostro et al., 2013) available at https://github.com/buenrostrolab/mitoblacklist/tree/master/fasta). Peak files for peaks with FDR < 0.05 per cell type and organ were intersected, keeping intervals called in at least two replicates. Resulting consensus lists were merged to create a final reference set of peaks. A matrix of raw counts as basis for differential analysis was created with featureCounts from the subread package. Only samples with a FRiP score above 0.05 were kept for downstream analysis.

For differential analysis, raw counts were normalized using the normOffsets function from the csaw package (Lun and Smyth, 2016). Regions were tested for differential accessibility per celltype between organs with the edgeR glmQLF framework, requiring a minimal absolute fold change beyond 1.5 (glmTreat function) at an FDR < 0.05. Differential regions per tested contrast were merged and clustered based on Z-scored logCPMs from edgeR using hierarchical clustering with ward.D2 agglomeration. Heatmaps were created with ComplexHeatmap.

Motif enrichments per cluster were obtained using the Analysis of Motif Enrichment (AME) algorithm from the meme suite (McLeay and Bailey, 2010) executed via the R/Bioconductor package memes (https://pubmed.ncbi.nlm.nih.gov/34570758/). Enrichment scores for significantly-enriched (FDR < 10e-5) motifs as basis for the heatmap in **Figure 4** were calculated as scaled-log10(X) where X represents the value as returned by AME.

### Neutrophil subpopulation isolation

Mouse blood, bone-marrow-derived or splenic immature and mature neutrophils were isolated from blood, tibias and femurs or the spleen respectively from male or female C57BL6/J mice, stained with fluorescently-labelled antibodies and sorted using a FACS Aria III (BD) or Sorter SH800S (Sony) based on Lineage^-^ (B220, CD3, F4/80, CD115), DAPI^-^, SSC^high^, CXCR2^low^ (immature) or CXCR2^high^ (mature).

### Neutrophil progenitor differentiation assay

Sorted preNEUs (Lineage^-^: B220, CD19, CD3, F4/80, CD115, TER119, Ly6C^+^, CD11b^+^, cKIT^med^, CXCR4^high^) were incubated in RPMI+1%FCS and recombinant GM-CSF (50 ng/mL), G-CSF (Neupogen, 50 ng/mL) or bone marrow supernatant. Bone marrow supernatant was obtained by flushing femur and tibia from C57BL6/J treated with CPM/AMD with RPMI+1%FCS. After the removal of cells by centrifugation for 5 minutes at 300G, supernatants were used at a 1:10 ratio with RPMI+1%FCS for preNEU differentiation. After 5 days of differentiation, descendant cells were quantified using flow cytometry.

### Analysis of reactive oxygen species

Blood, bone marrow or splenic mouse cells were incubated with 25 μM Dihidroetidium (DHE, Abcam) in RPMI containing 1% FCS for 10 min at 37°C. Directly after incubation, cells were incubated with LPS from *E.coli* K12 (1 ng/mL, Invivogen) for 30 minutes at 37°C. After washing, cells were incubated with fluorescent antibodies (CD45, Ly6G, CD11b, CD18, CXCR2, CD101) and DAPI. Upon oxidation, DHE is converted into its fluorescent form thereby acting as an indicator of the amount of reactive oxygen species present. The red fluorescence intensity generated by this dye was measured on immature or mature neutrophils using flow cytometry.

### Neutrophil extracellular trap analysis

Blood neutrophils (10000-80000 cells), bone marrow (50000 cells) or spleen (50000 cells) were seeded for 30 minutes at 37°C in HBSS + 25mM HEPES, followed by an incubation with LPS E.coli K12 (1 ng/mL, blood or 10 ng/mL, bone marrow and spleen) for 4 hours at 37°C After washing, neutrophils were fixed with 2% PFA and stained with anti-mouse citrullinated histone H3 (Abcam), anti-mouse Ly6G (clone 1A8, BD), anti-MPO (R&D) and DAPI. The immunofluorescence signal was measured using a fluorescent confocal microscope.

## Statistics

Statistical analysis was performed using GraphPad Prism 10 software (GraphPad Software). The ROUT outlier function was used to exclude statistical outliers (Q = 1%). Normal distribution of the data was assessed using the D’Agostino-Pearson omnibus test for normality. Normally distributed data was tested by two-tailed unpaired t-test (one variable) or one-way ANOVA with Tukey or Dunnet’s correction (>2 variables). When 2 factors were analyzed, data was analyzed using two-way ANOVA with Tukey’s correction. In all tests a 95% confidence interval was used, for which P < 0.05 was considered a significant difference. All data are represented as mean ± SEM. For in vivo experiments, we performed sample size calculation using https://homepage.stat.uiowa.edu/~rlenth/Power/. Animal were randomly distributed across experimental conditions and analysis was performed after blinding.

## Data and code availability

The data that support the findings of this study are available from the corresponding authors upon reasonable request.

## Declaration of generative AI and AI-assisted technologies

ChatGPT 4o was used during the generation and improvement of the R code used for scRNA-seq, mass-cytometry or spatial analysis. DeepL write was used to improve English grammar. After using this tool, the authors reviewed and edited the content as needed and take full responsibility for the content of the publication.

---

**Extended Figure 1.**
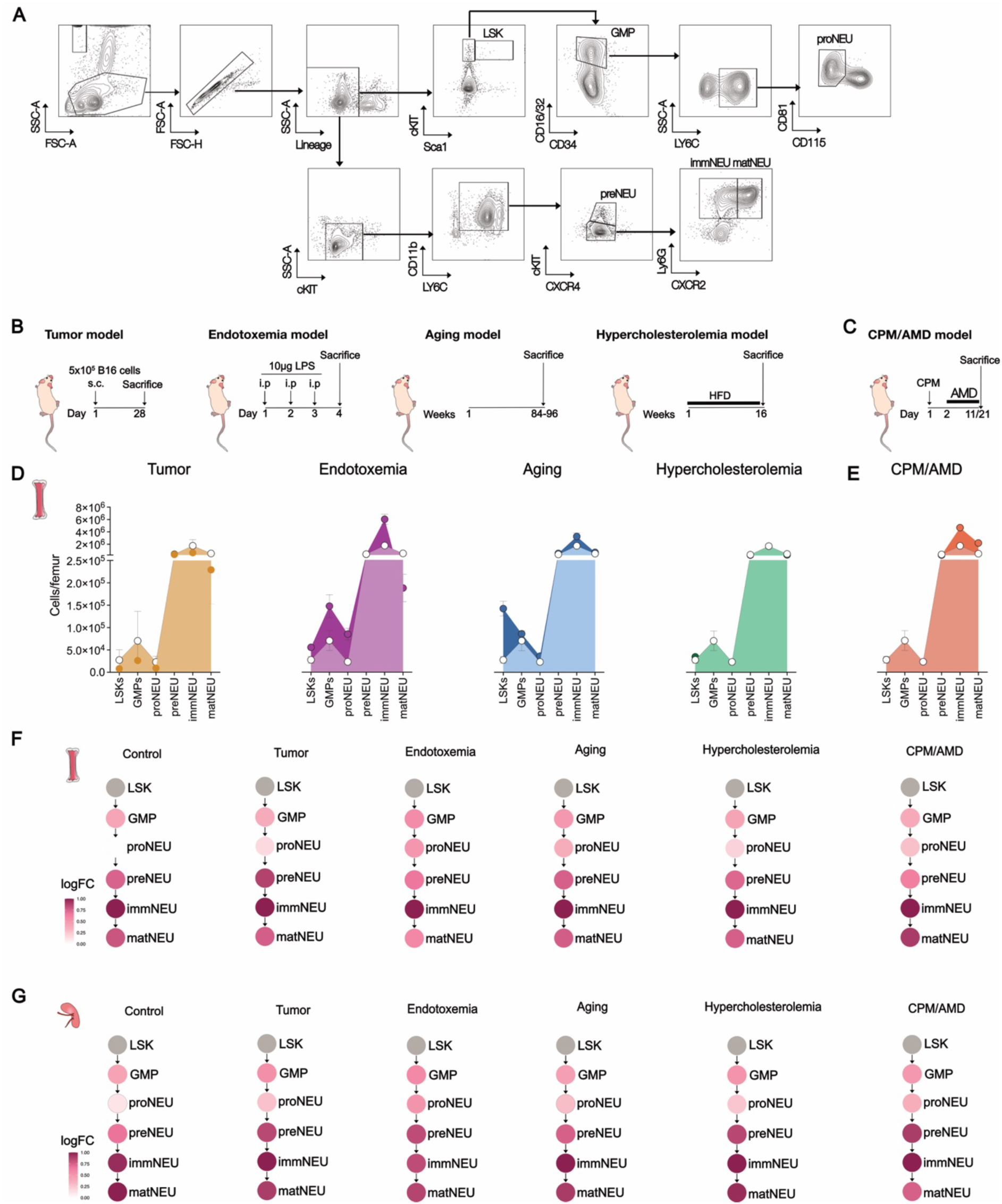
**Flow cytometry analysis of extramedullary granulopoiesis across pathological states**. **(A)** Flow cytometry gating strategy. **(B)** Scheme of mouse models: 8-12 weeks old female C57BL6/J mice. From left to right: tumor model of melanoma (50,000 B16 cells were injected subcutaneously in the left flank of the mouse and the tumor was retrieved 1 month later), chronic endotoxemia (daily injection of 10 μg of lipopolysaccharide intraperitoneally for 3 consecutive days), aging (84-96 weeks old mice), hypercholesterolemia (*Apoe^-/-^*female mice fed with high-cholesterol diet for 16 weeks) **(C)** CPM/AMD: C57BL6/J female mice were administered with cyclophosphamide (CPM), followed by daily administration of 5 mg/kg of AMD3100 (AMD). (**D/E**) Flow cytometry-based quantification of indicated stages of granulopoiesis in the bone marrow. Empty circles indicate respective bone marrow cell populations under steady-state (control) condition, filled circles represent counts of indicated maturation stage in the respective disease model. Analysis of proportions of each indicated subpopulation relative to LSKs (Lin^-^Sca1^+^cKit^+^) in bone marrow (**F**) and spleen (**G**). Granulocyte-macrophage progenitors (GMP, Lin^-^Sca1^-^ cKit^+^CD16/32^high^CD34^+^), neutrophil-specified (proNEUs, Lin^-^Sca1^-^cKit^+^ CD16/32^high^Ly6C^+^CD115^-^CD81^+^CD11b^+^) and-committed progenitors (preNEU, Lin^-^ cKit^int^Ly6C^+^CD115^-^CD11b^+^CXCR4^+^), immature (immNEU, CD11b^+^Ly6G^+^CD115^-^ CXCR2^-^) and mature neutrophils (matNEU, CD11b^+^Ly6G^+^CD115^-^CXCR2^+^). Data points indicate the mean of 3-10 mice. Data are mean +/-SEM.

**Extended Figure 2:**
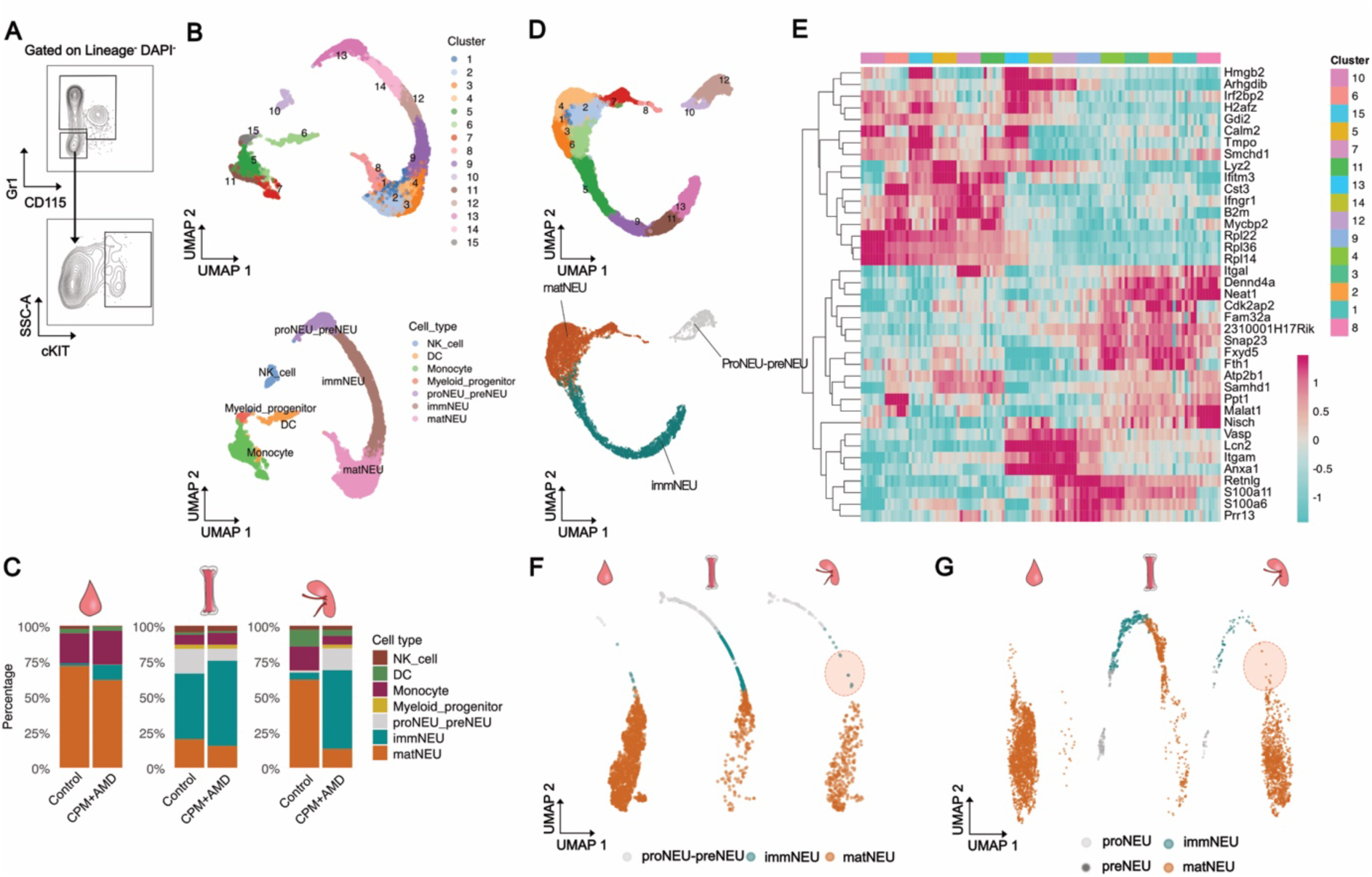
**scRNA-seq analysis of extramedullary granulopoiesis**. (**A**) Gating strategy for isolation of cells used for scRNA-seq analysis. **(B)** Uniform manifold approximation and projection (UMAP) plot of aggregated cells from blood, bone marrow and spleen colored by cluster (upper panel) and cell type (lower panel) from 4 mice per condition. **(C)** Proportion quantification from identified cell populations in scRNA-seq analysis. **(D)** UMAP plot of aggregated neutrophil progenitors and descendants from blood, bone marrow and spleen colored by cluster (upper panel) and cell type (lower panel). **(E)** Heatmap showing top expressed genes by cluster from (D). **(F/G)** UMAP of aggregated neutrophils from blood, bone marrow and spleen from hypercholesterolemic mice (*Apoe^-/-^* female mice fed with high-cholesterol diet for 16 weeks, a pool of 5 mice F) or tumor-bearing mice (C57BL6/J injected with 1 x 10^6^ in 100 µl PBS of murine oropharyngeal carcinoma subcutaneously, n = 4 mice per condition G) colored by cell type.

**Extended Figure 3.**
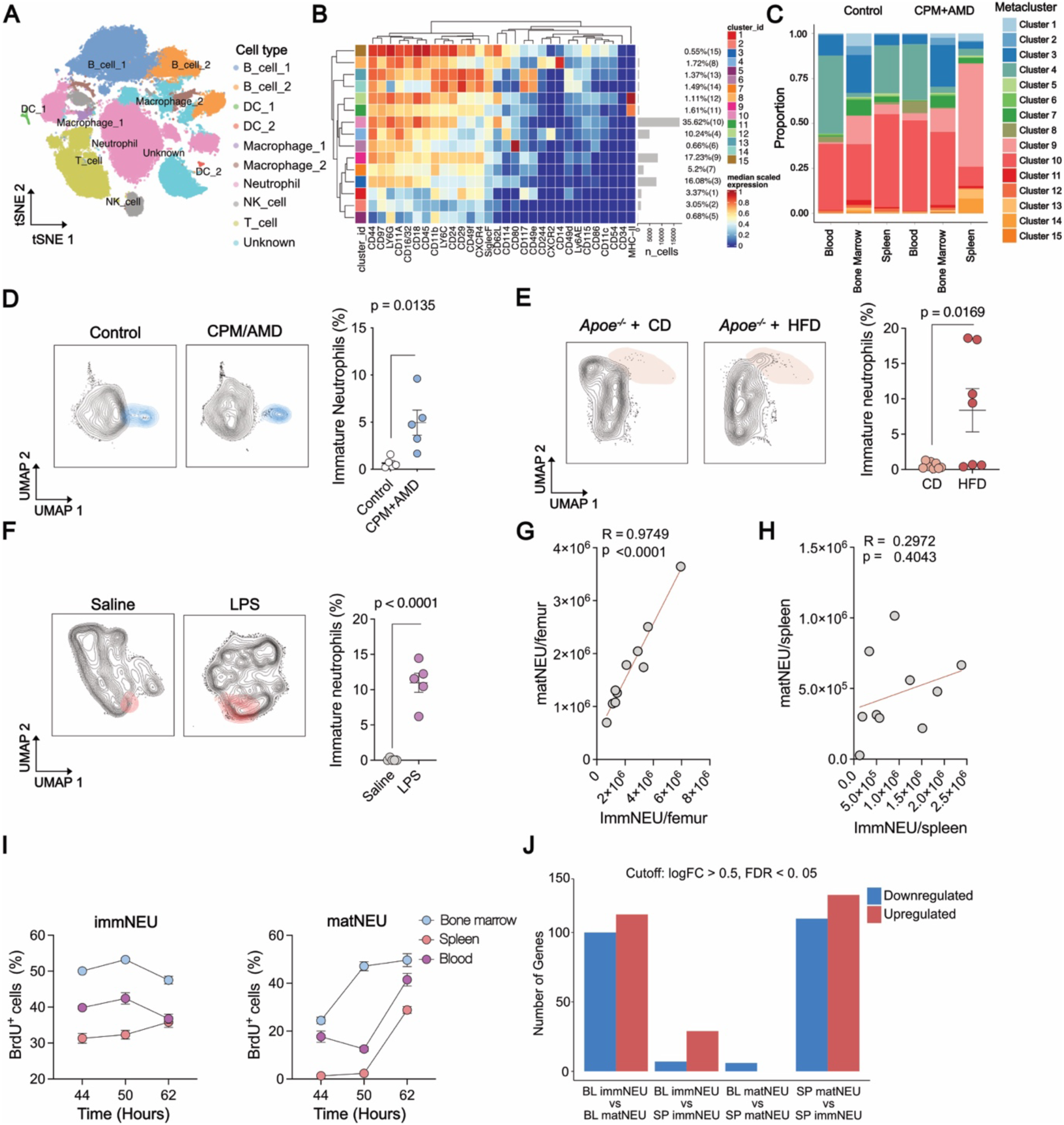
**Neutrophils of splenic origin are immature and mobilized to the blood in emergency granulopoiesis**. **(A-C)** C57BL6/J female mice were administered with cyclophosphamide (CPM), followed by daily administration of 5 mg/kg of AMD3100 (AMD) or saline for controls. Mass cytometry analysis of blood, bone marrow and spleen. **(A)** tSNE plot showing aggregated cells colored by cell type. **(B)** Heatmap showing scaled expression of surface markers grouped by metacluster. **(C)** Proportions of metaclusters by organ and treatment. **(D-F)** Quantification of circulating immature neutrophils by spectral flow cytometry in varying treatment conditions. Panels show UMAPs (left) depicting immature neutrophils in colored areas and the quantification of the percentage of circulating immature neutrophils (right). Unpaired t-test analysis. Data are mean +/-SEM. **(D)** C57BL6/J female mice were administered with CPM, followed by daily administration of 5 mg/kg of AMD or saline for controls. n = 5 mice/group. **(E)** Mouse model of hypercholesterolemia (*Apoe^-/-^* female mice fed with high-cholesterol diet (HFD) or chow diet (CD) for 16 weeks). n = 7-8 mice/group. **(F)** Mouse model of repeated LPS injection (10 μg/day i.p. for three days). n = 5 mice/group. **(G/H)** Pearson correlation between mature and immature neutrophils in the bone marrow (G) and spleen (H). **(I)** CPM/AMD treated mice were injected with a single dose of BrdU (100 μg) 44, 50 and 62 hours before sacrifice. Flow cytometric analysis of the percentage of BrdU^+^ immature (left panel, immNEU) and mature (right panel, matNEU) neutrophils in blood, bone marrow and spleen. n = 4 mice/group. Data are mean +/-SEM. **(J)** Number of differentially-expressed genes across indicated comparisons. BL: Blood; SP: Spleen.

**Extended Figure 4.**
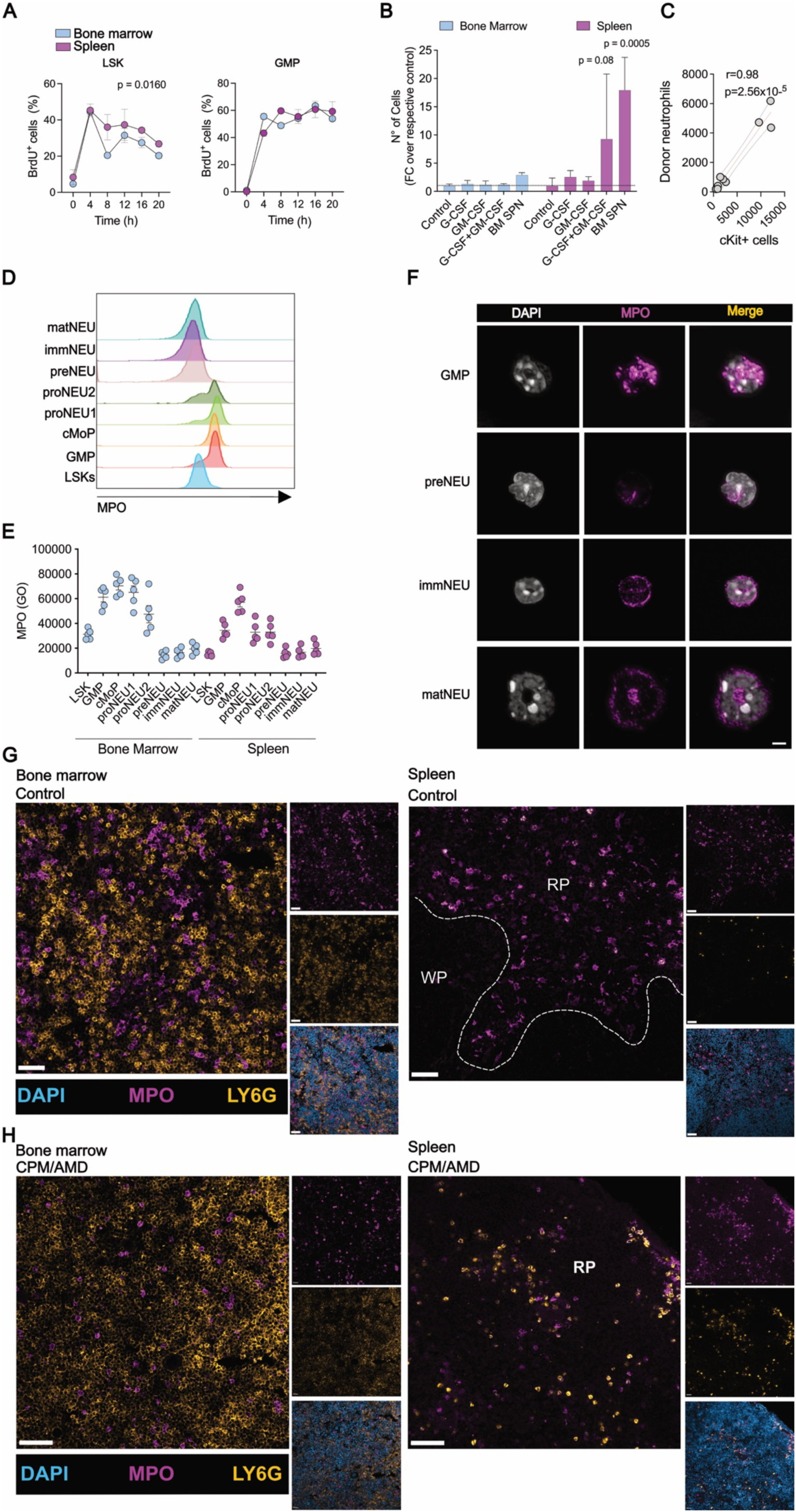
**Myeloperoxidase expression in myeloid progenitor and analysis of in vitro differentiation capacity**. C57BL6/J female mice were administered with cyclophosphamide (CPM), followed by daily administration of 5 mg/kg of AMD3100 (AMD) or saline for controls. **(A)** CPM/AMD-treated mice were injected with a pulse of 5‘Bromo-deoxyuridine (BrdU) and sacrificed at indicated time-points. Shown are the percentage of BrdU^+^ cells. Data are mean ± SEM. Two-way ANOVA with Šidák’s post-hoc correction. n = 2-4 mice/group. **(B)** Sorted preNEUs were incubated for 5 days with indicated treatments and descendant cells were enumerated using flow cytometry. G-CSF: Granulocyte-Colony Stimulating Factor (50 ng/mL); GM-CSF: Granulocyte Macrophage-Colony Stimulating Factor (50 ng/mL); BM SPN: Bone marrow supernatant; n = 2-3 mice/group. **(C)** Pearson correlation of donor neutrophils and donor cKit^+^ ells. **(D/E)** Flow cytometry analysis of myeloperoxidase (MPO) expression in indicated cell types across granulopoiesis. Histogram representation (C) and quantification (D) of mean fluorescence intensity. **(F)** Confocal microscopy images depicting MPO expression and nuclei (DAPI) on sorted granulocyte-macrophage progenitor (GMP), preNEU, immature (immNEU) and mature (matNEU) neutrophils. Scale bar: 4 μm. Data are mean ± SEM. Experiment performed once. (**G/H**) Immunofluorescence images depicting nuclei (DAPI, blue), Myeloperoxidase (MPO, magenta), Ly6G (ochre) in bone marrow and spleen. Images showing bone marrow and spleen in vehicle-(G) and CPM/AMD-(H) treated mice. RP: Red pulp; WP: White pulp. Scale bar: 100 μm.

**Extended Figure 5.**
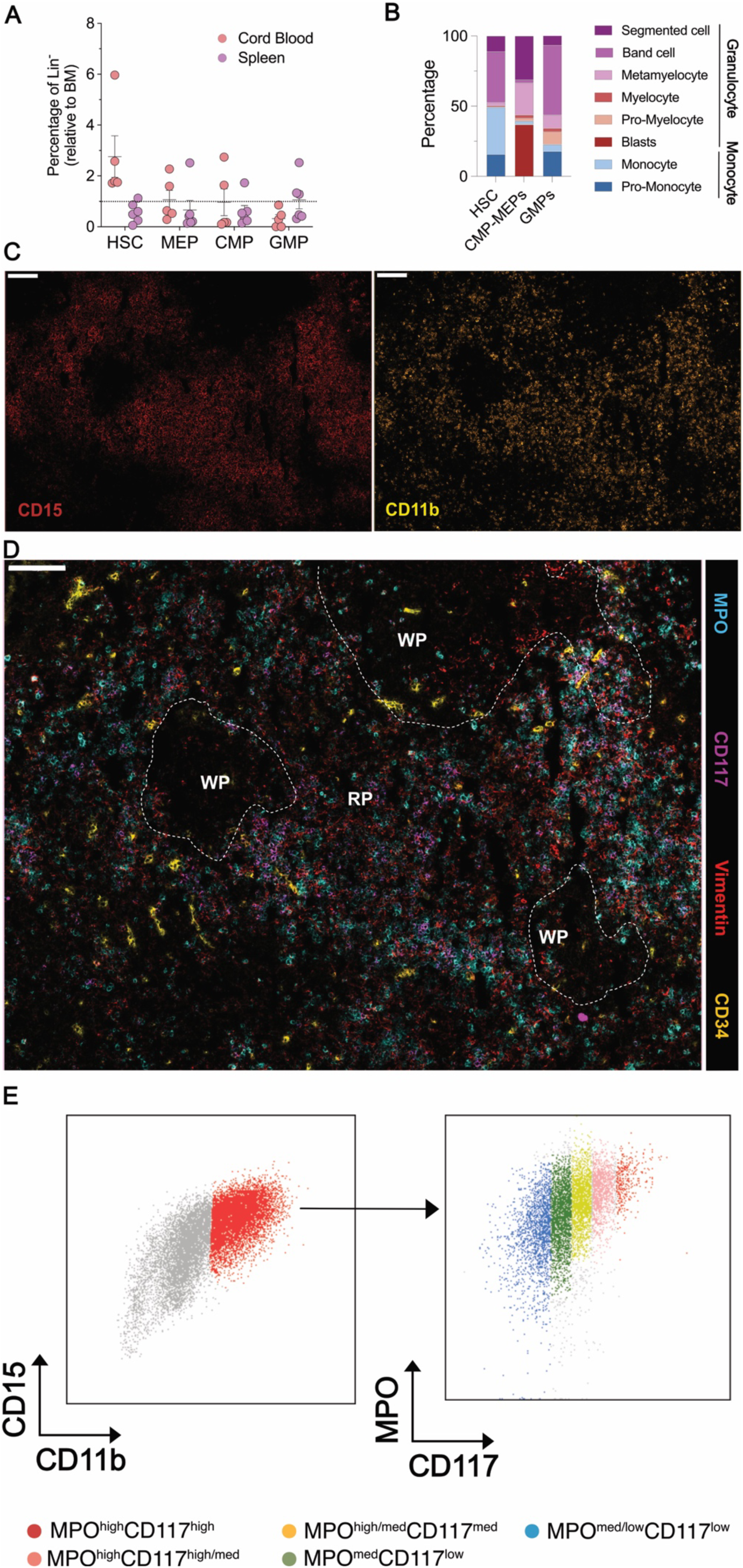
Human splenic granulopoiesis. **(A)** Flow cytometry analysis of human HSC (hematopoietic stem cell; lineage^-^CD45^med^CD34^+^), MEP (myeloid-erythroid progenitor; lineage^-^CD45^med^CD34^+^CD38^+^CD45R^-^IL3Ra^-^), CMP (common myeloid progenitor; lineage^-^CD45^med^CD34^+^CD38^+^CD45R^-^IL3Ra^+^), GMP (granulocyte-macrophage progenitor; lineage^-^CD45^med^CD34^+^CD38^+^CD45R^+^IL3Ra^+^) in samples of bone marrow, blood and spleen. Data are mean ± SEM. n = 5-6 donors/group. **(B)** Stacked bar graph showing the percentage of descendants of spleen progenitors (Pro-Monocyte: CD15^dim/-^CD16^-^CD64^bright^ CD14^dim/-^; Monocyte: CD15^dim/-^CD16^-^ CD64^+^CD14^bright^; Pro-Myelocyte: CD15^dim^CD16^-^CD64^+^CD14^-^SSC^int^; Myelocyte: CD15^dim^CD16^-^CD64^+^CD14^-^SSC^high^; Metamyelocyte: CD15^+^CD16^dim^CD64^dim^ CD14^-^ CD10^-^; Band cell: CD15^+^CD16^+^CD64^-^CD14^-^CD10^+^; Segmented cell: CD15^bright^CD16^bright^CD64^-^CD14^-^CD10^+^) analyzed by flow cytometry 14 days after differentiation. n = 1-3 donors/group (**C**) Immunofluorescence images depicting CD15 (red, left image) and CD11b (ochre, right image) from human spleen specimens. Scale bar: 100 μm. (**D**) Immunofluorescence images depicting MPO (cyan), CD117 (magenta), Vimentin (red) and CD34 (ochre) from human spleen specimens. Dashed lines delineate the white pulp (WP). RP: Red pulp. Scale bar: 100 μm. (**E**) Gating strategy for the identification of MPO^high^CD117^high^ (red), MPO^high^CD117^high/med^ (pink), MPO^high^CD117^med^ (orange), MPO^med^CD117^low^ (green), MPO^low^CD117^low^ (blue) cells.

**Extended Figure 6.**
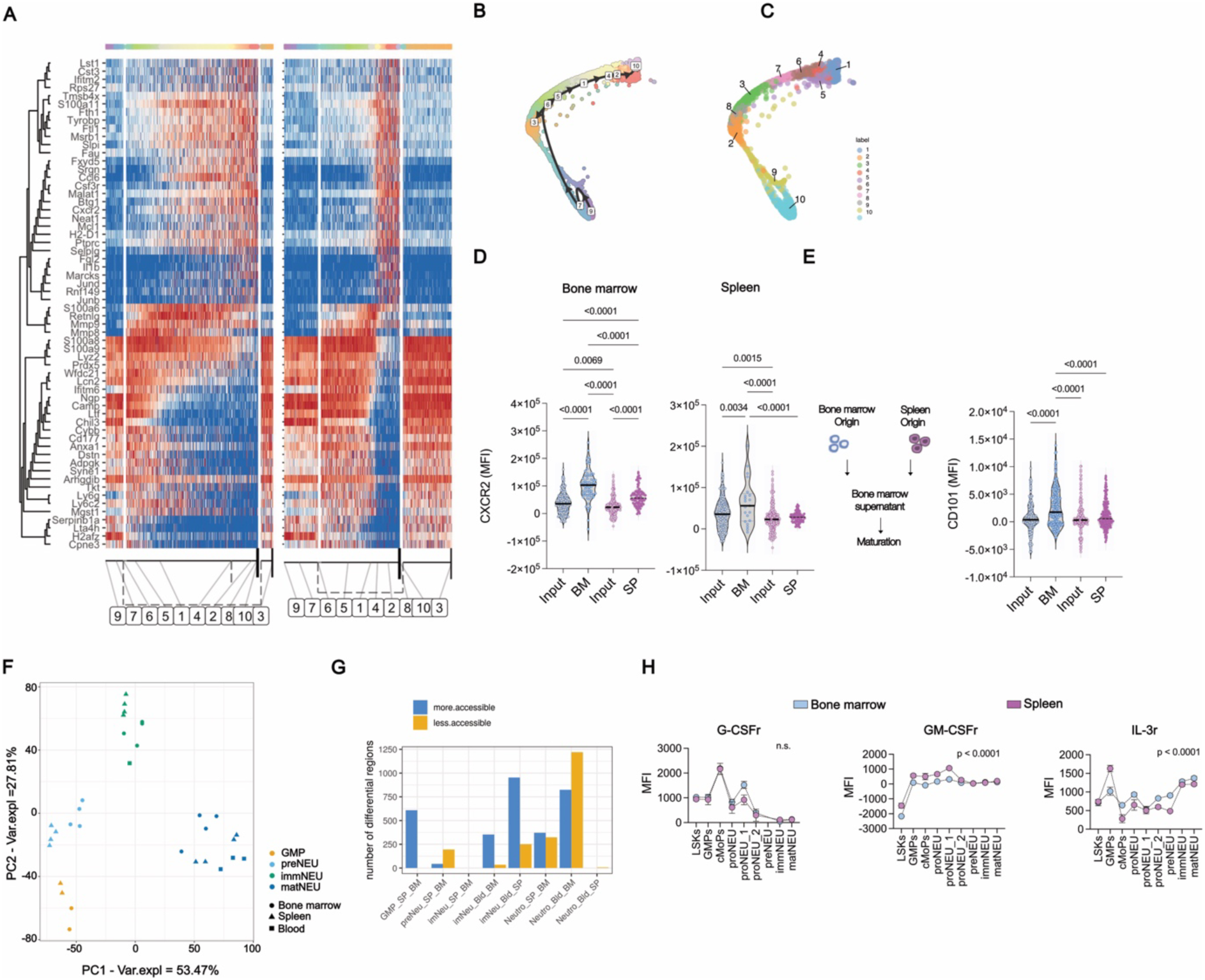
sNeuPs show increased open chromatin sites and surface-activated phenotype to generate immature neutrophils with defective maturation. C57BL6/J female mice were administered with cyclophosphamide (CPM), followed by daily administration of 5 mg/kg of AMD3100 (AMD) or saline for controls. **(A)** Heatmap showing scaled expression of genes defining identified PAGA trajectories and represented by milestones ranges. Milestones represent the dynamic transition between cell states. PAGA trajectory analysis on scRNA-seq of aggregated neutrophils from blood, bone marrow and spleen colored by clusters defined by milestone-enriched cells **(B)** or Louvain clustering **(C)**. Immature neutrophils (B220^-^ CD19^-^CD3^-^CD115^-^F4/80^-^SSC^high^CXCR2^low^) isolated from bone marrow (dsRED) or spleen (Lyz2-eGFP) were adoptively transferred intravenously into control mice **(D)** or incubated with bone marrow supernatant for 12 hours **(E).** MFI: Mean fluorescent intensity. **(D)** Flow cytometry analysis of CXCR2 expression of donor neutrophils isolated from the recipient bone marrow or spleen. Data is represented respectively to CXCR2 levels in the input. n = 20-200 cells out of 6 mice. MFI: Mean fluorescent intensity. **(E)** Flow cytometry analysis of CD101 expression of neutrophils after in vitro maturation. Data is represented respective to CD101 levels in the input. n = 180-198 cells out of 6 mice. Representative results from 1 out of two independent experiments. One-way ANOVA analysis with Tukey’s correction. ****P<0.0001. MFI: Mean fluorescent intensity. **(F)** PCA analysis on indicated cell populations from ATACseq analysis. **(G)** Bar graph showing the number of differential open chromatin regions for indicated comparisons. SP: Spleen; BM: Bone marrow; Bld: Blood. **(H)** Flow cytometry analysis of surface expression of Granulocyte-Colony Stimulating Factor receptor (G-CSFr, left), Granulocyte Macrophage-Colony Stimulating Factor receptor (GM-CSFr, middle) and Interleukin-3 receptor (IL-3r, right) of indicated cells. Two-way ANOVA analysis. n = 4-5 mice. Data are mean +/-SEM.

**Extended Figure 7.**
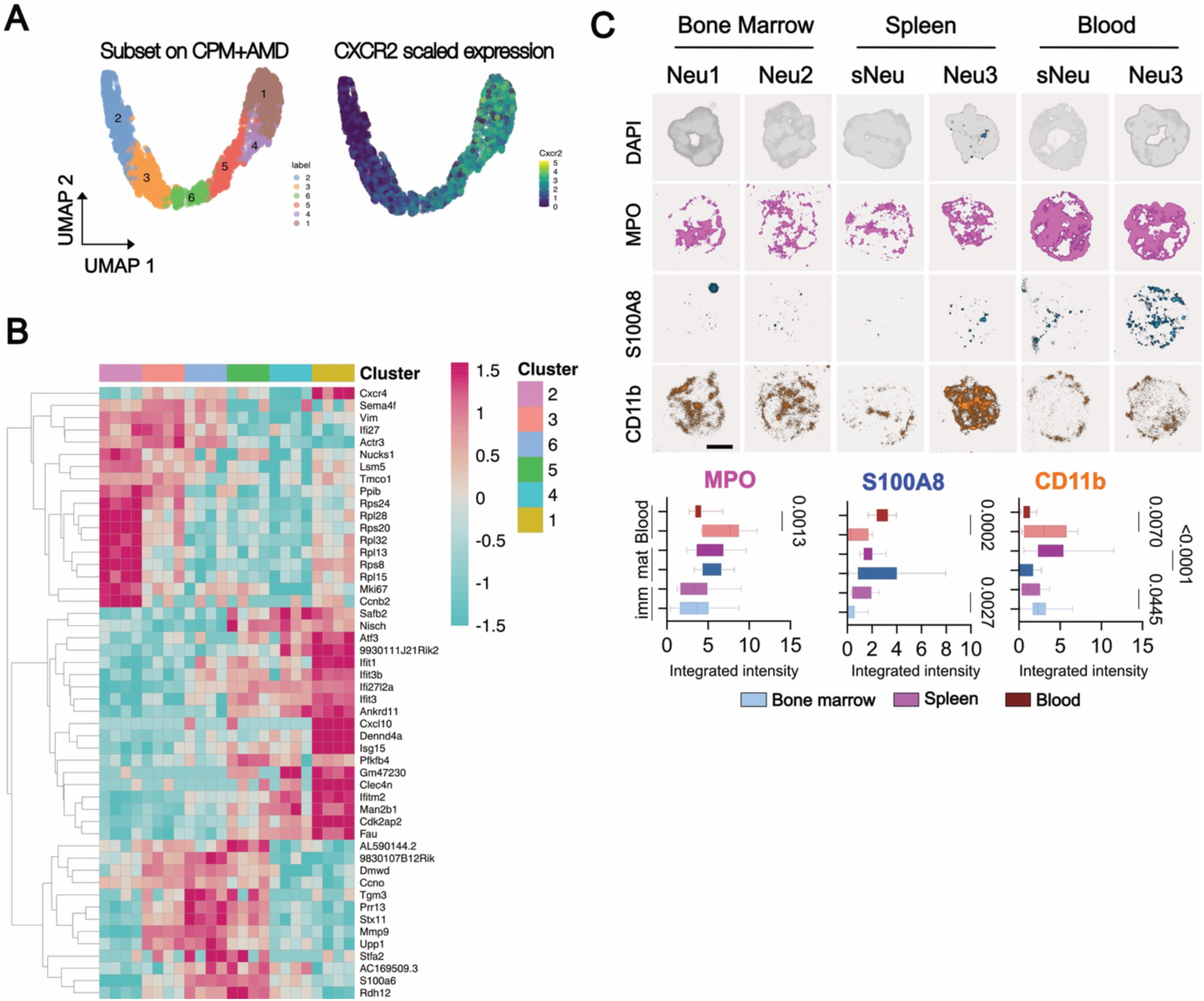
**Circulating sNeu exhibit increased myeloperoxidase and CD11b expression**. C57BL6/J female mice were administered with cyclophosphamide (CPM), followed by daily administration of 5 mg/kg of AMD3100 (AMD) or saline for controls. **(A/B)** scRNA-seq on bone marrow ad splenic neutrophils from CPM/AMD model. **(A)** UMAP showing unsupervised clustering (left) and scaled Cxcr2 expression indicating maturation direction (right). **(B)** Heatmap showing top expressed genes by cluster from (A). (**C**) Immunofluorescence on FACS-sorted mature (B220^-^CD19^-^CD3^-^CD115^-^F4/80^-^SSC^high^CXCR2^high^) or immature (B220^-^CD19^-^ CD3^-^CD115^-^F4/80^-^SSC^high^CXCR2^low^) from blood, bone marrow or spleen showing nuclei (DAPI, gray), Myeloperoxidase (MPO, magenta), S100A8 (blue) and CD11b (orange). N = 7-51 cells from 2-4 mice. Scale bar: 4 μm. One-way ANOVA with Tukey’s post-hoc correction. Data are mean +/-SEM.

**Extended Figure 8.**
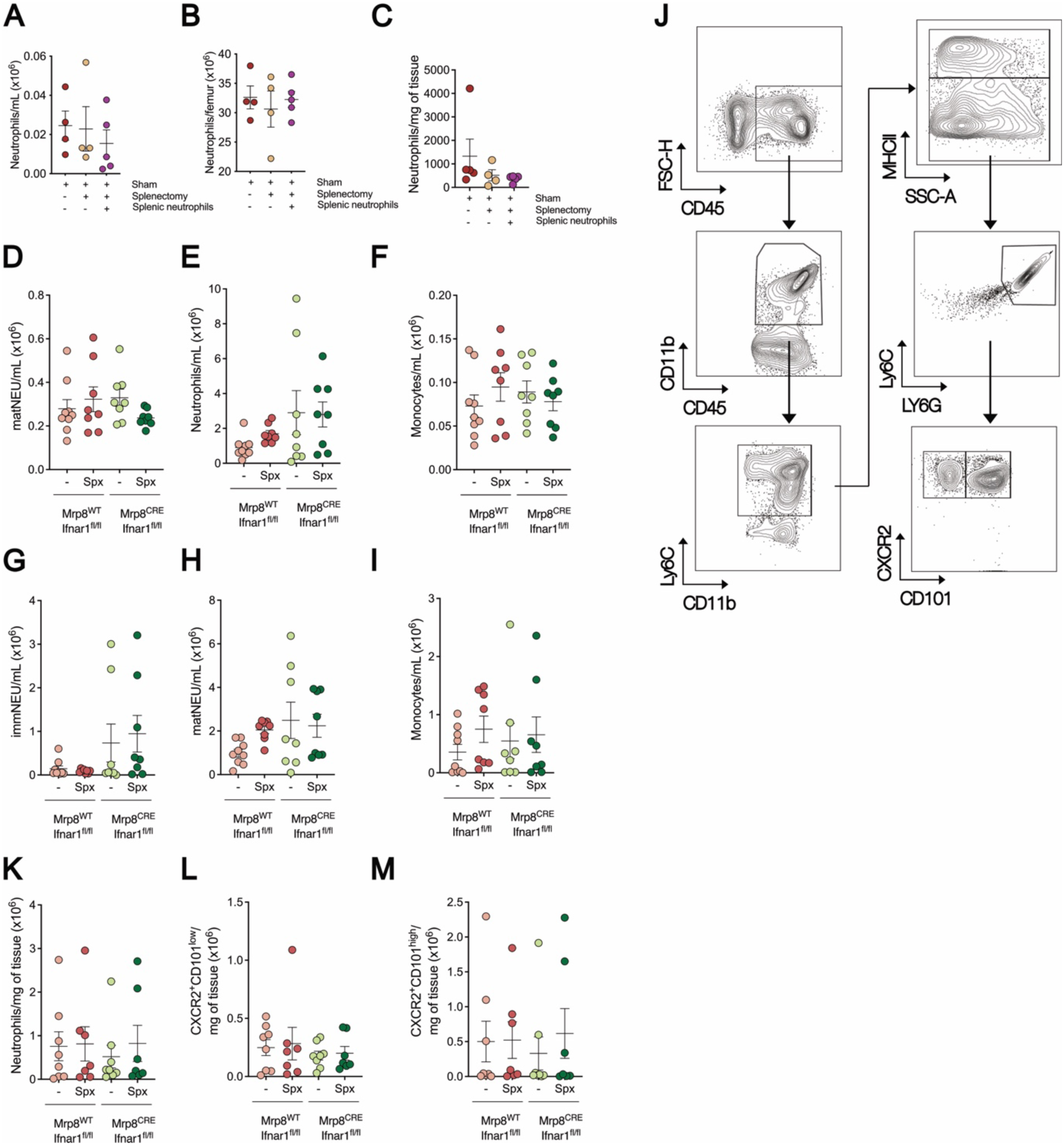
**Impact of splenectomy on leukocyte subpopulation in CPM/AMD-treated mice after bacterial infection**. **(A-C)** Mice were subjected to splenectomy (Spx) or sham operation. A group of splenectomized mice was administered with splenic neutrophils. All groups were infected with 5×10^8^ uropathogenic *E.coli* and sacrifice 2 hours post-infection. Flow cytometry analysis of circulating (A), bone marrow (B) or tissue bladder (C) neutrophils. **(D-M)** Mice were lethally irradiated and transplanted with bone marrow cells from *Mrp8^cre^Ifnar1^fl/fl^*or *Mrp8^wt^Ifnar1^fl/fl^*. After reconstitution, mice were subjected to splenectomy or sham, followed by treatment with CPM/AMD. Flow cytometry analysis of mature neutrophils (D), total neutrophils (E) and monocytes (F) in blood pre-infection. Flow cytometry analysis of immature neutrophils (G), mature neutrophils (H) and monocytes (I) in blood post-infection. (J) Flow cytometry gating strategy used in tissue bladder. Flow cytometry analysis of total (K), immature (L), and mature (M) neutrophils. Results are represented as mean±SEM.

## Supplementary Material

**Table 1.**
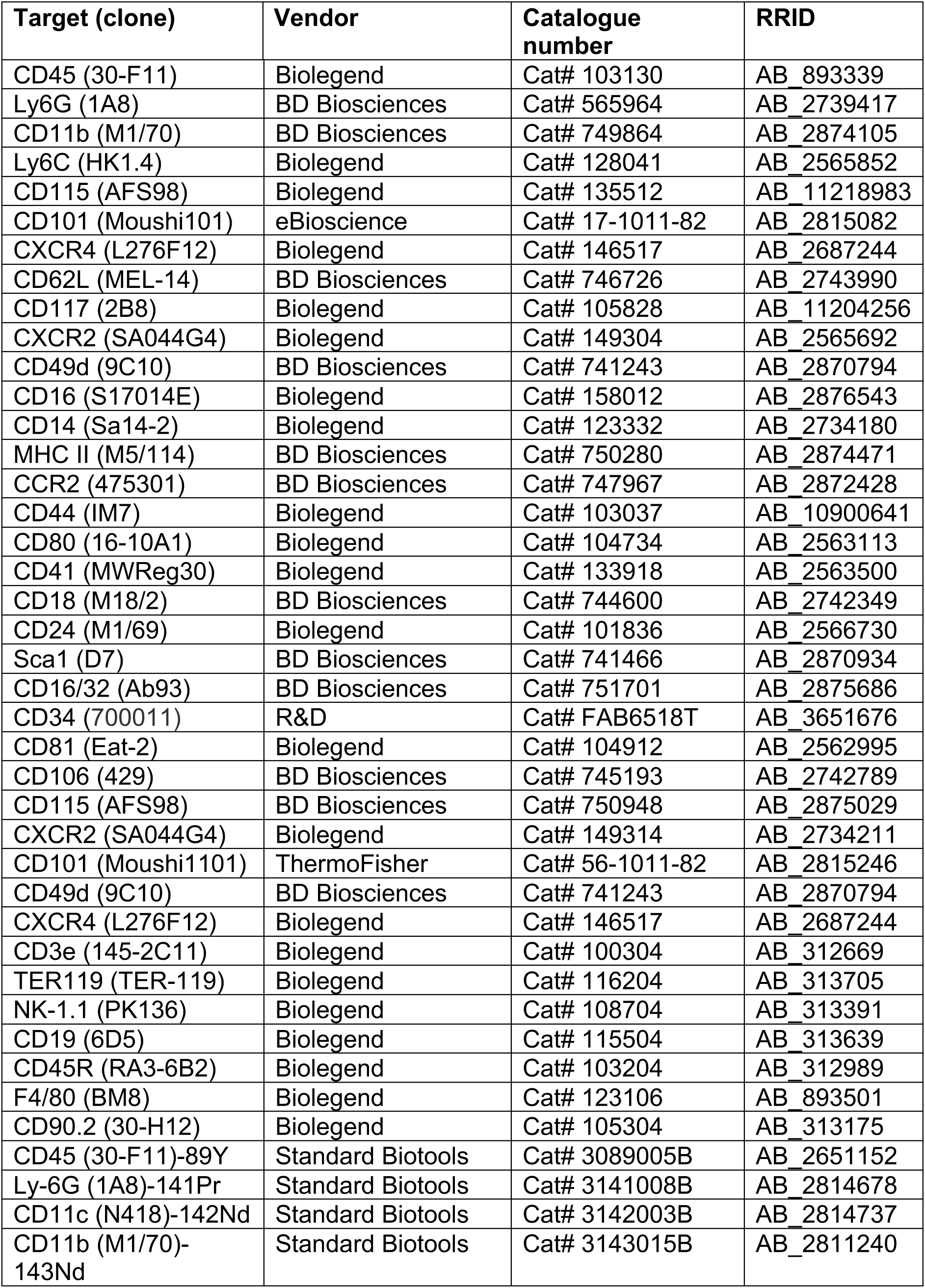
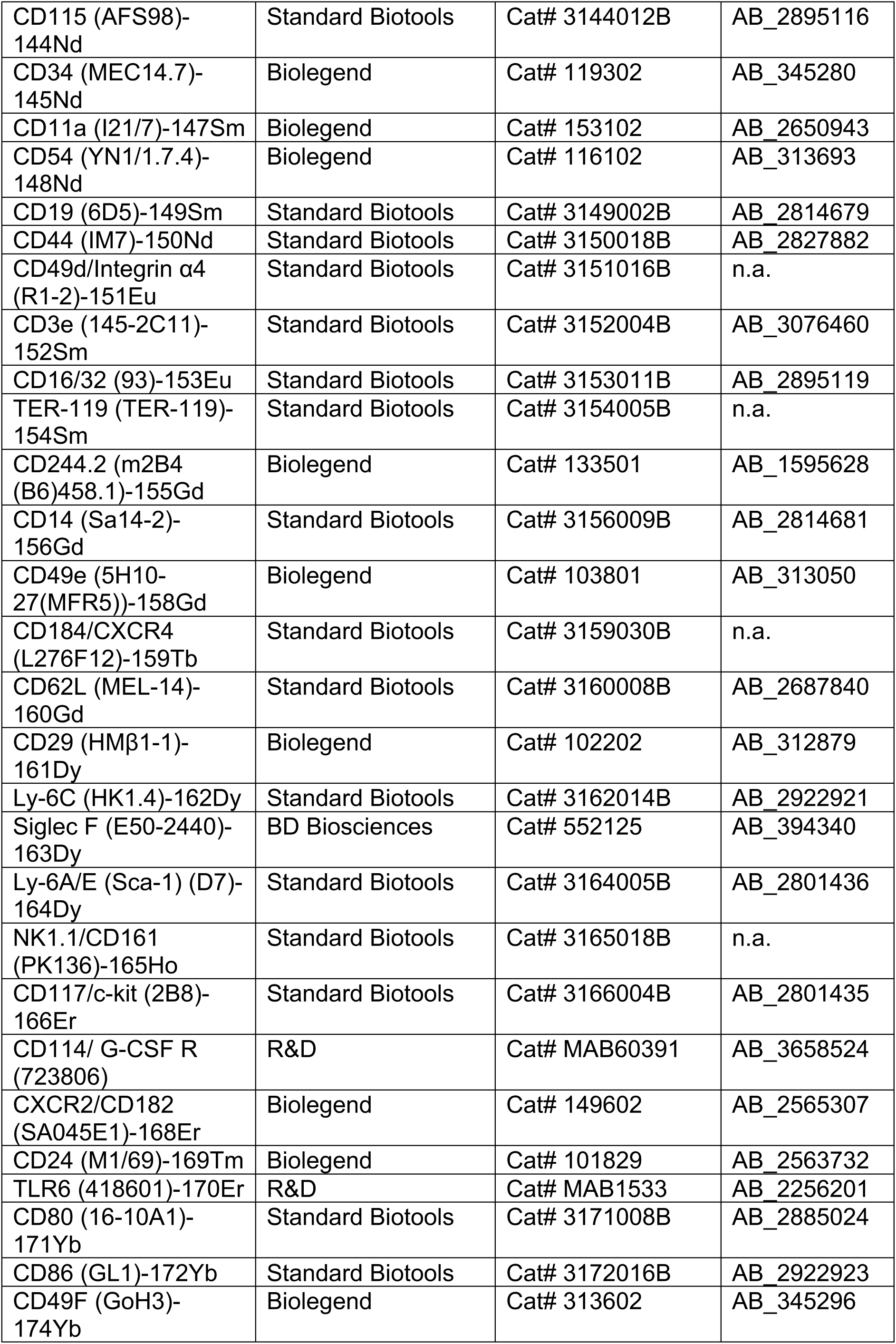
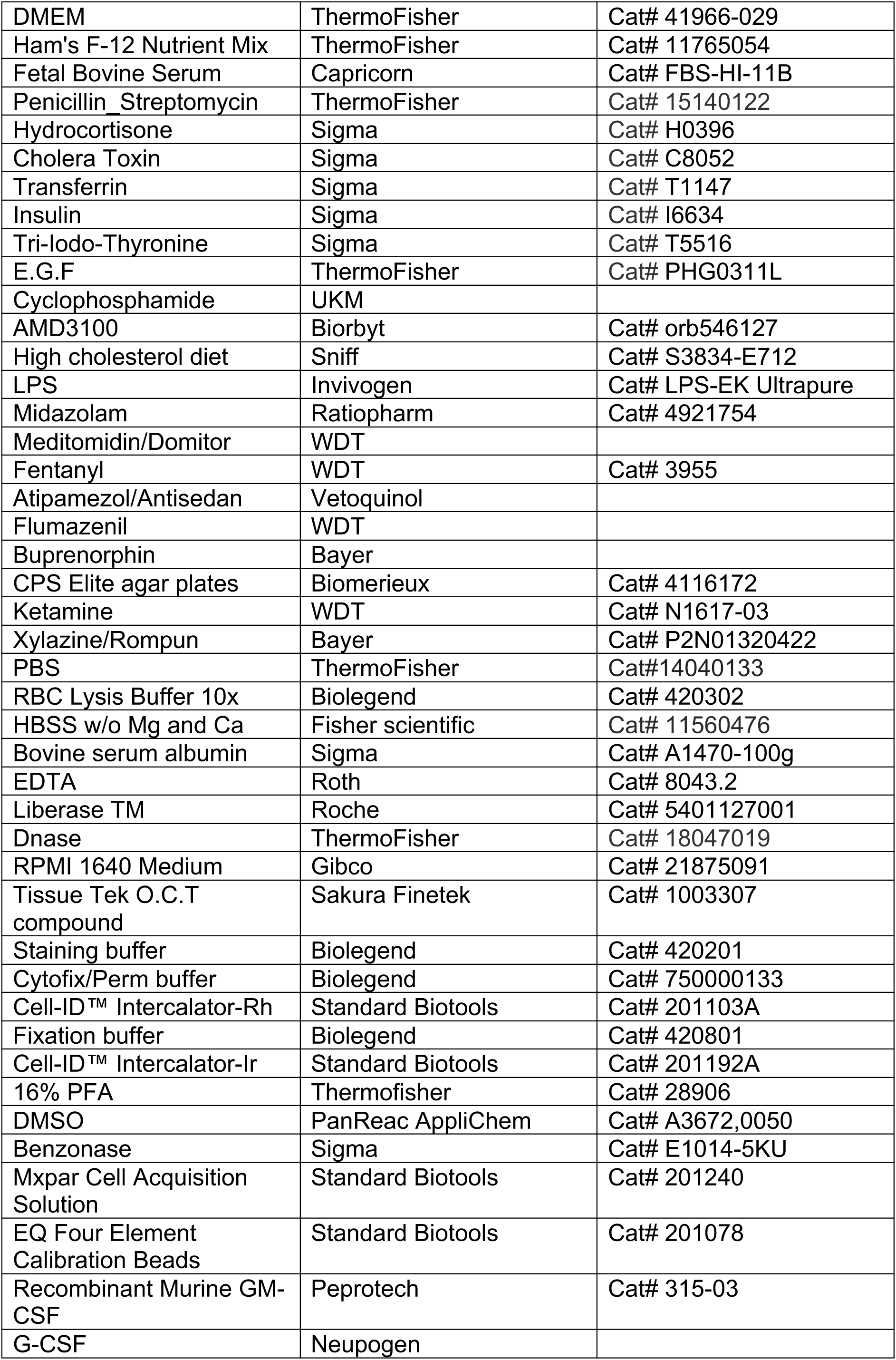
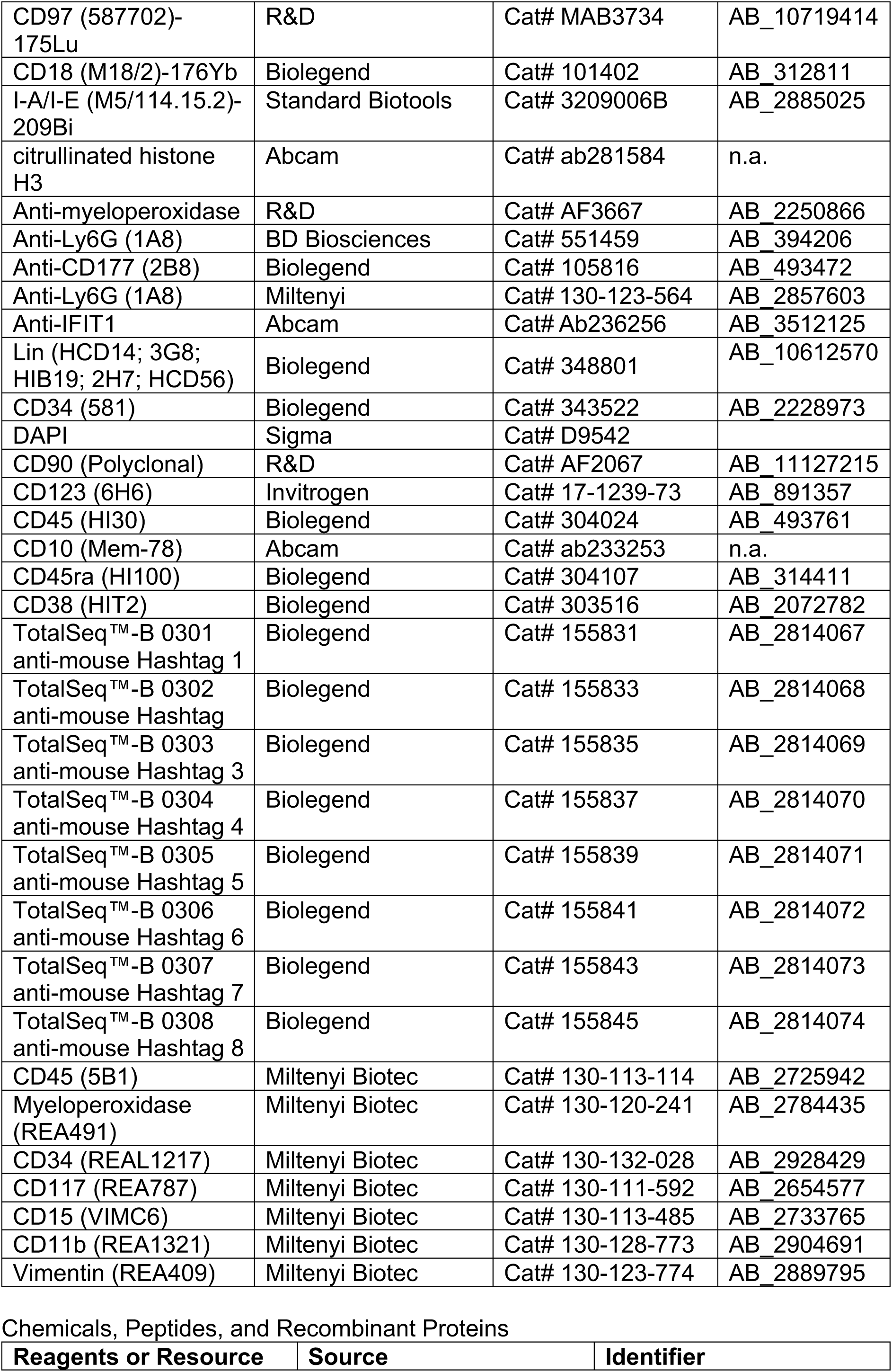
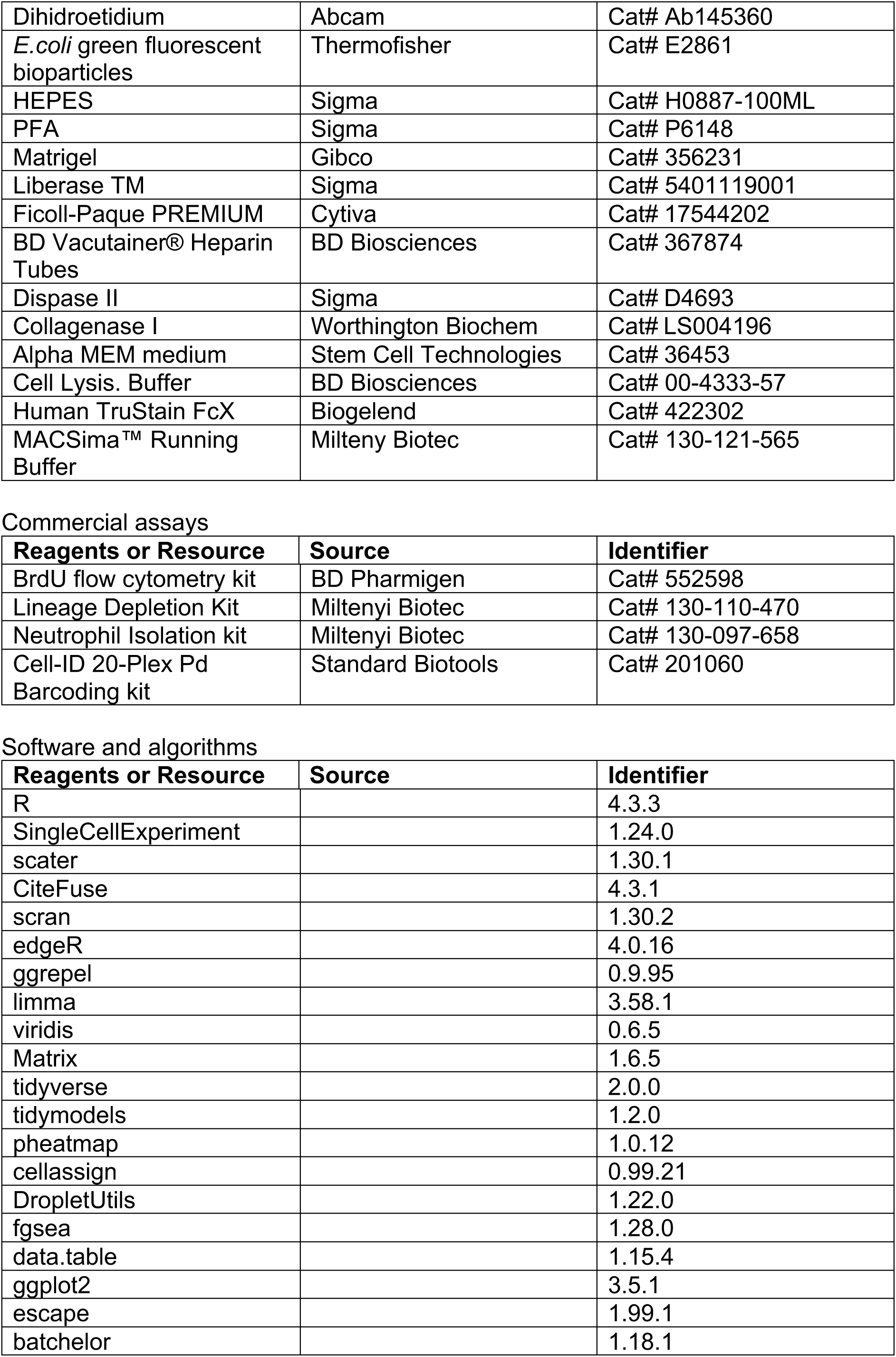
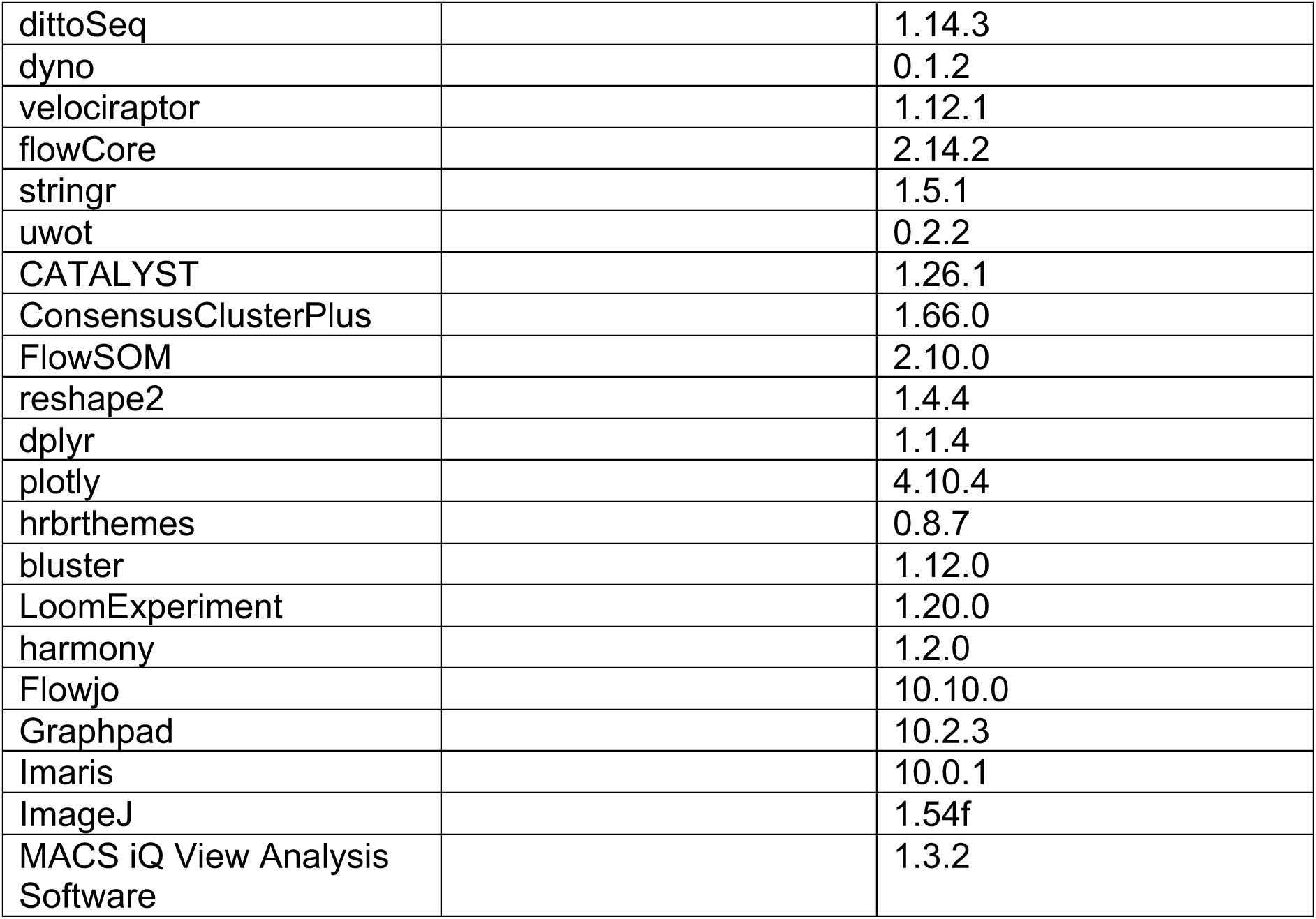
Antibodies.

## References

Adrover, J.M., Aroca-Crevillen, A., Crainiciuc, G., Ostos, F., Rojas-Vega, Y., Rubio-Ponce, A., Cilloniz, C., Bonzon-Kulichenko, E., Calvo, E., Rico, D., et al. (2020). Programmed’disarming’ of the neutrophil proteome reduces the magnitude of inflammation. Nat Immunol 21, 135–144.

Adrover, J.M., Del Fresno, C., Crainiciuc, G., Cuartero, M.I., Casanova-Acebes, M., Weiss, L.A., Huerga-Encabo, H., Silvestre-Roig, C., Rossaint, J., Cossio, I., et al. (2019). A Neutrophil Timer Coordinates Immune Defense and Vascular Protection. Immunity 50, 390–402 e310.

Amemiya, H.M., Kundaje, A., and Boyle, A.P. (2019). The ENCODE Blacklist: Identification of Problematic Regions of the Genome. Sci Rep 9, 9354.

Andreatta, M., and Carmona, S.J. (2021). UCell: Robust and scalable single-cell gene signature scoring. Comput Struct Biotechnol J 19, 3796–3798.

Ballesteros, I., Rubio-Ponce, A., Genua, M., Lusito, E., Kwok, I., Fernandez-Calvo, G., Khoyratty, T.E., van Grinsven, E., Gonzalez-Hernandez, S., Nicolas-Avila, J.A., et al. (2020). Co-option of Neutrophil Fates by Tissue Environments. Cell 183, 1282–1297 e1218.

Bayne, L.J., Beatty, G.L., Jhala, N., Clark, C.E., Rhim, A.D., Stanger, B.Z., and Vonderheide, R.H. (2012). Tumor-derived granulocyte-macrophage colony-stimulating factor regulates myeloid inflammation and T cell immunity in pancreatic cancer. Cancer Cell 21, 822–835.

Bergen, V., Lange, M., Peidli, S., Wolf, F.A., and Theis, F.J. (2020). Generalizing RNA velocity to transient cell states through dynamical modeling. Nat Biotechnol 38, 1408–1414.

Borcherding, N., Vishwakarma, A., Voigt, A.P., Bellizzi, A., Kaplan, J., Nepple, K., Salem, A.K., Jenkins, R.W., Zakharia, Y., and Zhang, W. (2021). Mapping the immune environment in clear cell renal carcinoma by single-cell genomics. Commun Biol 4, 122.

Buenrostro, J.D., Giresi, P.G., Zaba, L.C., Chang, H.Y., and Greenleaf, W.J. (2013). Transposition of native chromatin for fast and sensitive epigenomic profiling of open chromatin, DNA-binding proteins and nucleosome position. Nat Methods 10, 1213–1218.

Burberry, A., Zeng, M.Y., Ding, L., Wicks, I., Inohara, N., Morrison, S.J., and Nunez, G. (2014). Infection mobilizes hematopoietic stem cells through cooperative NOD-like receptor and Toll-like receptor signaling. Cell Host Microbe 15, 779–791.

Carissimo, G., Xu, W., Kwok, I., Abdad, M.Y., Chan, Y.H., Fong, S.W., Puan, K.J., Lee, C.Y., Yeo, N.K., Amrun, S.N., et al. (2020). Whole blood immunophenotyping uncovers immature neutrophil-to-VD2 T-cell ratio as an early marker for severe COVID-19. Nat Commun 11, 5243.

Casanova-Acebes, M., Nicolas-Avila, J.A., Li, J.L., Garcia-Silva, S., Balachander, A., Rubio-Ponce, A., Weiss, L.A., Adrover, J.M., Burrows, K., N, A.G., et al. (2018). Neutrophils instruct homeostatic and pathological states in naive tissues. J Exp Med 215, 2778–2795.

Casanova-Acebes, M., Pitaval, C., Weiss, L.A., Nombela-Arrieta, C., Chevre, R., N, A.G., Kunisaki, Y., Zhang, D., van Rooijen, N., Silberstein, L.E., et al. (2013). Rhythmic modulation of the hematopoietic niche through neutrophil clearance. Cell 153, 1025–1035.

Coffelt, S.B., Kersten, K., Doornebal, C.W., Weiden, J., Vrijland, K., Hau, C.S., Verstegen, N.J.M., Ciampricotti, M., Hawinkels, L., Jonkers, J., et al. (2015). IL-17-producing gammadelta T cells and neutrophils conspire to promote breast cancer metastasis. Nature 522, 345–348.

Coppin, E., Florentin, J., Vasamsetti, S.B., Arunkumar, A., Sembrat, J., Rojas, M., and Dutta, P. (2018). Splenic hematopoietic stem cells display a pre-activated phenotype. Immunol Cell Biol.

Corces, M.R., Trevino, A.E., Hamilton, E.G., Greenside, P.G., Sinnott-Armstrong, N.A., Vesuna, S., Satpathy, A.T., Rubin, A.J., Montine, K.S., Wu, B., et al. (2017). An improved ATAC-seq protocol reduces background and enables interrogation of frozen tissues. Nat Methods 14, 959–962.

Cortez-Retamozo, V., Etzrodt, M., Newton, A., Rauch, P.J., Chudnovskiy, A., Berger, C., Ryan, R.J., Iwamoto, Y., Marinelli, B., Gorbatov, R., et al. (2012). Origins of tumor-associated macrophages and neutrophils. Proc Natl Acad Sci U S A 109, 2491–2496.

Drifte, G., Dunn-Siegrist, I., Tissieres, P., and Pugin, J. (2013). Innate immune functions of immature neutrophils in patients with sepsis and severe systemic inflammatory response syndrome. Crit Care Med 41, 820–832.

Dutta, P., Hoyer, F.F., Grigoryeva, L.S., Sager, H.B., Leuschner, F., Courties, G., Borodovsky, A., Novobrantseva, T., Ruda, V.M., Fitzgerald, K., et al. (2015). Macrophages retain hematopoietic stem cells in the spleen via VCAM-1. J Exp Med 212, 497–512.

Emami, H., Singh, P., MacNabb, M., Vucic, E., Lavender, Z., Rudd, J.H., Fayad, Z.A., Lehrer-Graiwer, J., Korsgren, M., Figueroa, A.L., et al. (2015). Splenic metabolic activity predicts risk of future cardiovascular events: demonstration of a cardiosplenic axis in humans. JACC Cardiovasc Imaging 8, 121–130.

Evrard, M., Kwok, I.W.H., Chong, S.Z., Teng, K.W.W., Becht, E., Chen, J., Sieow, J.L., Penny, H.L., Ching, G.C., Devi, S., et al. (2018). Developmental Analysis of Bone Marrow Neutrophils Reveals Populations Specialized in Expansion, Trafficking, and Effector Functions. Immunity 48, 364–379 e368.

Herault, A., Binnewies, M., Leong, S., Calero-Nieto, F.J., Zhang, S.Y., Kang, Y.A., Wang, X., Pietras, E.M., Chu, S.H., Barry-Holson, K., et al. (2017). Myeloid progenitor cluster formation drives emergency and leukaemic myelopoiesis. Nature 544, 53–58.

Hsu, B.E., Tabaries, S., Johnson, R.M., Andrzejewski, S., Senecal, J., Lehuede, C., Annis, M.G., Ma, E.H., Vols, S., Ramsay, L., et al. (2019). Immature Low-Density Neutrophils Exhibit Metabolic Flexibility that Facilitates Breast Cancer Liver Metastasis. Cell Rep 27, 3902–3915 e3906.

Hussain, T., Domnich, M., Bordbari, S., Pylaeva, E., Siakaeva, E., Spyra, I., Ozel, I., Droege, F., Squire, A., Lienenklaus, S., et al. (2022). IFNAR1 Deficiency Impairs Immunostimulatory Properties of Neutrophils in Tumor-Draining Lymph Nodes. Front Immunol 13, 878959.

Inra, C.N., Zhou, B.O., Acar, M., Murphy, M.M., Richardson, J., Zhao, Z., and Morrison, S.J. (2015). A perisinusoidal niche for extramedullary haematopoiesis in the spleen. Nature 527, 466–471.

Khoyratty, T.E., Ai, Z., Ballesteros, I., Eames, H.L., Mathie, S., Martin-Salamanca, S., Wang, L., Hemmings, A., Willemsen, N., von Werz, V., et al. (2021). Distinct transcription factor networks control neutrophil-driven inflammation. Nat Immunol 22, 1093–1106.

Kim, E.J., Kim, S., Kang, D.O., and Seo, H.S. (2014). Metabolic activity of the spleen and bone marrow in patients with acute myocardial infarction evaluated by 18f-fluorodeoxyglucose positron emission tomograpic imaging. Circ Cardiovasc Imaging 7, 454–460.

Klein, J.C., Moses, K., Zelinskyy, G., Sody, S., Buer, J., Lang, S., Helfrich, I., Dittmer, U., Kirschning, C.J., and Brandau, S. (2017). Combined toll-like receptor 3/7/9 deficiency on host cells results in T-cell-dependent control of tumour growth. Nat Commun 8, 14600.

Kwok, I., Becht, E., Xia, Y., Ng, M., Teh, Y.C., Tan, L., Evrard, M., Li, J.L.Y., Tran, H.T.N., Tan, Y., et al. (2020). Combinatorial Single-Cell Analyses of Granulocyte-Monocyte Progenitor Heterogeneity Reveals an Early Uni-potent Neutrophil Progenitor. Immunity 53, 303–318 e305.

Leuschner, F., Rauch, P.J., Ueno, T., Gorbatov, R., Marinelli, B., Lee, W.W., Dutta, P., Wei, Y., Robbins, C., Iwamoto, Y., et al. (2012). Rapid monocyte kinetics in acute myocardial infarction are sustained by extramedullary monocytopoiesis. J Exp Med 209, 123–137.

Li, X., Wang, H., Yu, X., Saha, G., Kalafati, L., Ioannidis, C., Mitroulis, I., Netea, M.G., Chavakis, T., and Hajishengallis, G. (2022). Maladaptive innate immune training of myelopoiesis links inflammatory comorbidities. Cell 185, 1709–1727 e1718.

Lun, A.T., and Smyth, G.K. (2016). csaw: a Bioconductor package for differential binding analysis of ChIP-seq data using sliding windows. Nucleic Acids Res 44, e45.

Mackey, J.B.G., Coffelt, S.B., and Carlin, L.M. (2019). Neutrophil Maturity in Cancer. Front Immunol 10, 1912.

Magri, G., Miyajima, M., Bascones, S., Mortha, A., Puga, I., Cassis, L., Barra, C.M., Comerma, L., Chudnovskiy, A., Gentile, M., et al. (2014). Innate lymphoid cells integrate stromal and immunological signals to enhance antibody production by splenic marginal zone B cells. Nat Immunol 15, 354–364.

Manz, M.G., and Boettcher, S. (2014). Emergency granulopoiesis. Nat Rev Immunol 14, 302–314.

McLeay, R.C., and Bailey, T.L. (2010). Motif Enrichment Analysis: a unified framework and an evaluation on ChIP data. BMC Bioinformatics 11, 165.

Mende, N., and Laurenti, E. (2021). Hematopoietic stem and progenitor cells outside the bone marrow: where, when, and why. Exp Hematol 104, 9–16.

Moon, K.R., van Dijk, D., Wang, Z., Gigante, S., Burkhardt, D.B., Chen, W.S., Yim, K., Elzen, A.V.D., Hirn, M.J., Coifman, R.R., et al. (2019). Visualizing structure and transitions in high-dimensional biological data. Nat Biotechnol 37, 1482–1492.

Muench, D.E., Olsson, A., Ferchen, K., Pham, G., Serafin, R.A., Chutipongtanate, S., Dwivedi, P., Song, B., Hay, S., Chetal, K., et al. (2020). Mouse models of neutropenia reveal progenitor-stage-specific defects. Nature 582, 109–114.

Ohtsu, S., Yagi, H., Nakamura, M., Ishii, T., Kayaba, S., Soga, H., Gotoh, T., Rikimaru, A., Kokubun, S., and Itoh, T. (2000). Enhanced neutrophilic granulopoiesis in rheumatoid arthritis. Involvement of neutrophils in disease progression. J Rheumatol 27, 1341–1351.

Prada-Medina, C.A., Peron, J.P.S., and Nakaya, H.I. (2020). Immature neutrophil signature associated with the sexual dimorphism of systemic juvenile idiopathic arthritis. J Leukoc Biol 108, 1319–1327.

Pylaeva, E., Korschunow, G., Spyra, I., Bordbari, S., Siakaeva, E., Ozel, I., Domnich, M., Squire, A., Hasenberg, A., Thangavelu, K., et al. (2022). During early stages of cancer, neutrophils initiate anti-tumor immune responses in tumor-draining lymph nodes. Cell Rep 40, 111171.

Regan-Komito, D., Swann, J.W., Demetriou, P., Cohen, E.S., Horwood, N.J., Sansom, S.N., and Griseri, T. (2020). GM-CSF drives dysregulated hematopoietic stem cell activity and pathogenic extramedullary myelopoiesis in experimental spondyloarthritis. Nat Commun 11, 155.

Rice, C.M., Lewis, P., Ponce-Garcia, F.M., Gibbs, W., Groves, S., Cela, D., Hamilton, F., Arnold, D., Hyams, C., Oliver, E., et al. (2023). Hyperactive immature state and differential CXCR2 expression of neutrophils in severe COVID-19. Life Sci Alliance 6.

Robbins, C.S., Chudnovskiy, A., Rauch, P.J., Figueiredo, J.L., Iwamoto, Y., Gorbatov, R., Etzrodt, M., Weber, G.F., Ueno, T., van Rooijen, N., et al. (2012). Extramedullary hematopoiesis generates Ly-6C(high) monocytes that infiltrate atherosclerotic lesions. Circulation 125, 364–374.

Saelens, W., Cannoodt, R., Todorov, H., and Saeys, Y. (2019). A comparison of single-cell trajectory inference methods. Nat Biotechnol 37, 547–554.

Schaupp, L., Muth, S., Rogell, L., Kofoed-Branzk, M., Melchior, F., Lienenklaus, S., Ganal-Vonarburg, S.C., Klein, M., Guendel, F., Hain, T., et al. (2020). Microbiota-Induced Type I Interferons Instruct a Poised Basal State of Dendritic Cells. Cell 181, 1080–1096 e1019.

Schmidt, U., Weigert, M., Broaddus, C., and Myers, E.W. (2018). Cell Detection with Star-convex Polygons. Paper presented at: International Conference on Medical Image Computing and Computer-Assisted Intervention.

Schulte-Schrepping, J., Reusch, N., Paclik, D., Bassler, K., Schlickeiser, S., Zhang, B., Kramer, B., Krammer, T., Brumhard, S., Bonaguro, L., et al. (2020). Severe COVID-19 Is Marked by a Dysregulated Myeloid Cell Compartment. Cell 182, 1419–1440 e1423.

Silvestre-Roig, C., Braster, Q., Ortega-Gomez, A., and Soehnlein, O. (2020). Neutrophils as regulators of cardiovascular inflammation. Nat Rev Cardiol 17, 327–340.

Silvestre-Roig, C., Fridlender, Z.G., Glogauer, M., and Scapini, P. (2019). Neutrophil Diversity in Health and Disease. Trends Immunol 40, 565–583.

Silvin, A., Chapuis, N., Dunsmore, G., Goubet, A.G., Dubuisson, A., Derosa, L., Almire, C., Henon, C., Kosmider, O., Droin, N., et al. (2020). Elevated Calprotectin and Abnormal Myeloid Cell Subsets Discriminate Severe from Mild COVID-19. Cell 182, 1401–1418 e1418.

Swann, J.W., Olson, O.C., and Passegue, E. (2024). Made to order: emergency myelopoiesis and demand-adapted innate immune cell production. Nat Rev Immunol.

van Grinsven, E., Textor, J., Hustin, L.S.P., Wolf, K., Koenderman, L., and Vrisekoop, N. (2019). Immature Neutrophils Released in Acute Inflammation Exhibit Efficient Migration despite Incomplete Segmentation of the Nucleus. J Immunol 202, 207–217.

Wan, M., Lu, Y., Mao, B., Yu, S., Ju, P., Hu, K., Xu, Y., Li, X., and Zhuang, J. (2023). Immature neutrophil is associated with coronary plaque vulnerability based on optical coherence tomography analysis. Int J Cardiol 374, 89–93.

Weber, G.F., Chousterman, B.G., He, S., Fenn, A.M., Nairz, M., Anzai, A., Brenner, T., Uhle, F., Iwamoto, Y., Robbins, C.S., et al. (2015). Interleukin-3 amplifies acute inflammation and is a potential therapeutic target in sepsis. Science 347, 1260–1265.

Williams, R., Lee, D.W., Elzey, B.D., Anderson, M.E., Hostager, B.S., and Lee, J.H. (2009). Preclinical models of HPV+ and HPV-HNSCC in mice: an immune clearance of HPV+ HNSCC. Head Neck 31, 911–918.

Wu, C., Ning, H., Liu, M., Lin, J., Luo, S., Zhu, W., Xu, J., Wu, W.C., Liang, J., Shao, C.K., et al. (2018). Spleen mediates a distinct hematopoietic progenitor response supporting tumor-promoting myelopoiesis. J Clin Invest 128, 3425–3438.

Wu, Q., Zhang, J., Kumar, S., Shen, S., Kincaid, M., Johnson, C.B., Zhang, Y.S., Turcotte, R., Alt, C., Ito, K., et al. (2024). Resilient anatomy and local plasticity of naive and stress haematopoiesis. Nature 627, 839–846.

Xie, X., Shi, Q., Wu, P., Zhang, X., Kambara, H., Su, J., Yu, H., Park, S.Y., Guo, R., Ren, Q., et al. (2020). Single-cell transcriptome profiling reveals neutrophil heterogeneity in homeostasis and infection. Nat Immunol 21, 1119–1133.

Yvan-Charvet, L., and Ng, L.G. (2019). Granulopoiesis and Neutrophil Homeostasis: A Metabolic, Daily Balancing Act. Trends Immunol 40, 598–612.

Zhang, J., Wu, Q., Johnson, C.B., Pham, G., Kinder, J.M., Olsson, A., Slaughter, A., May, M., Weinhaus, B., D’Alessandro, A., et al. (2021). In situ mapping identifies distinct vascular niches for myelopoiesis. Nature 590, 457–462.

Zhu, Y.P., Padgett, L., Dinh, H.Q., Marcovecchio, P., Blatchley, A., Wu, R., Ehinger, E., Kim, C., Mikulski, Z., Seumois, G., et al. (2018). Identification of an Early Unipotent Neutrophil Progenitor with Pro-tumoral Activity in Mouse and Human Bone Marrow. Cell Rep 24, 2329–2341 e2328.

